# Mitophagy upregulates WNT5A/Ca^2+^ signalling to accelerate fibroblast migration and wound healing

**DOI:** 10.64898/2025.12.03.692038

**Authors:** Matthew Hunt, Monica Torres, Nuoqi Wang, Shannon Hinch, Margarita Chatzopoulou, Gustavo Urbano-Quispe, Etty Bachar-Wikström, Jakob D Wikström

**Author notes:** Corresponding author: Jakob Wikström, Dermato-Venereology Unit, Department of Molecular medicine (Solna), Karolinska Institutet, Sweden. The authors have declared that no conflict of interest exists.

## Abstract

In the event of dysregulated wound healing, hard-to-heal chronic wounds form and can place a significant burden on healthcare systems, yet gaps in knowledge surrounding the cellular and molecular processes involved have resulted in a lack of effective treatments. Here, we show that ubiquitin-independent mitophagy is upregulated in the early- and mid-wound healing stages. Additionally, enhancing mitophagy through Urolithin A treatment improved wound healing, in particular by accelerating fibroblast migration. RNAseq analysis demonstrated an upregulation of non-canonical WNT5A signalling in Urolithin A-treated fibroblasts, which was underpinned by elevated cytosolic Ca^2+^ buffering and CREB phosphorylation, ultimately leading to enhanced actin polymerisation and fibroblast migration. This study is thus the first to elucidate a role for mitophagy specifically in fibroblasts during wound healing; to demonstrate an important role for mitophagy in Ca^2+^-mediated WNT5A signalling cascades; and indicate the potential therapeutic benefits of treating non-healing wounds with mitophagy inducers such as Urolithin A.

## Introduction

Chronic wounds (CWs) place an enormous burden on healthcare systems, accounting for approximately 2-4% of healthcare budgets worldwide^1^, and are associated with a significant decrease in both morbidity and quality of life^2^. Although the pathophysiology of chronic wounds differs between patients, the most common causes are venous insufficiency (50-60%), arterial insufficiency (15-20%), and diabetes mellitus (5%)^3^.

Wound healing is a conserved process consisting of four concurrent and overlapping phases: haemostasis, inflammation, proliferation, and tissue remodelling^4^. CWs can arise in the instance of any of these phases becoming abhorrent, such as sustained inflammation, local tissue hypoxia, or impairment of cell motility during the proliferation stage^3^. Dermal fibroblasts are pleiomorphic and play vital roles in all stages of wound healing, including the deposition of extracellular matrix (ECM) components, wound contraction, and scar tissue formation^5,6^.

Importantly, wound healing is a highly metabolically demanding process and previous studies from our group and others have linked metabolic shifts towards glycolysis with improved fibroblast migration^7^, macrophage function^8^, neo-angiogenesis^9^, and accelerated wound healing in general^7^. Mitochondrial autophagy (mitophagy) is the selective sequestration and clearance of mitochondria by autophagy, and effective regulation of mitophagy is essential for both the functional integrity of the mitochondrial network and cell homeostasis^10,11^.

Indeed, defective mitophagy and the subsequent accumulation of dysfunctional mitochondria are implicated in several diseases^12^, and can be triggered by several stressors such as hypoxia, starvation, or mitochondrial damage^13^.

Currently, the precise roles of mitophagy in wound healing are incompletely understood^14^. Few studies have directly investigated the mechanistic role of mitophagy in skin wound healing, whilst the significance of mitophagy in fibroblast function during wound healing remains unexplored. However, of relevance, it has been demonstrated that mitophagy promoted keratinocyte migration and proliferation through the degradation of p-MAP4^15^ and separately, and that pharmacological induction of mitophagy promoted angiogenesis and collagen deposition in wound healing^16^. Additionally, it was shown in mouse models within non-wound healing contexts that BNIP3-induced mitophagy under the control of HIF-1α accelerated keratinocyte migration^17,18^, and that both BNIP3- and NIX-mediated mitophagy play important roles in keratinocyte differentiation in the epidermis^19–21^.

Herein, in this study we show that mitophagy is upregulated during normal wound healing, peaking at the proliferation stage, and that mitophagy is elevated in migrating fibroblasts *in vitro.* Pharmacological induction of mitophagy with urolithin A (UroA) increased wound closure in aged mouse models as well *ex vivo* full-thickness models, and increased fibroblast migration *in vitro*. Through transcriptomic and functional analysis in fibroblasts, we demonstrated that UroA-induced mitophagy resulted in the elevation of cytosolic Ca^2+^, leading to the phosphorylation of CREB and increased transcription of *WNT5A*. Elevated WNT5A protein levels led to the decreased capping of F-actin as well as increased actin polymerisation, ultimately resulting in accelerated fibroblast migration.

## Results

### Ub-independent mitophagy is upregulated in human wound healing samples

Initially, skin biopsies harvested from intact, 1 day post wounding (dpw), and 7 dpw healthy volunteer tissue, were subjected to H&E histochemistry in order to ensure samples were representative of classical wound healing (**Figures 1A and 1B**). Next, using publicly available RNA-sequencing (RNAseq) gene expression data obtained from a previous study^22^ – which utilised similar skin biopsies from healthy patients intact, 1 and 7 dpw wounded tissue, as well as venous ulcer (VU) CW tissue – we compared the differential gene expression (DEG) of various mitophagy and mitochondrial-function related genes between the respective groups (**Figures 1C and 1D**). Interestingly, we found that several ub-independent mitophagy-related genes were significantly differentially expressed in wounded skin compared to intact skin. Here, *BNIP3* expression was significantly upregulated in both 1 and 7 dpw as well as CW *vs* intact skin, whilst *NIX* and *FUNDC1* expression was significantly upregulated in 7 dpw *vs* intact skin. *BNIP3, NIX, PRKN,* and *OPTN* were all significantly higher in 7 dpw *vs* 1 dpw. Additionally, *DNM1L* expression was significantly upregulated in 1 and 7 dpw *vs* intact skin, and *MFN2* was significantly downregulated in 7 dpw *vs* intact skin. Finally, *BNIP3* expression was also significantly upregulated in CW *vs* intact skin, and both *BNIP3* and *NIX* expression was significantly higher in CW *vs* 1 dpw, but not compared to 7 dpw. Overall, RNAseq data suggested that ub-independant mitophagy pathways are upregulated in the early and mid-stages of wound healing, peaking at the proliferation stage, and that this coincides with increased mitochondrial fission gene expression.

**Figure 1.**
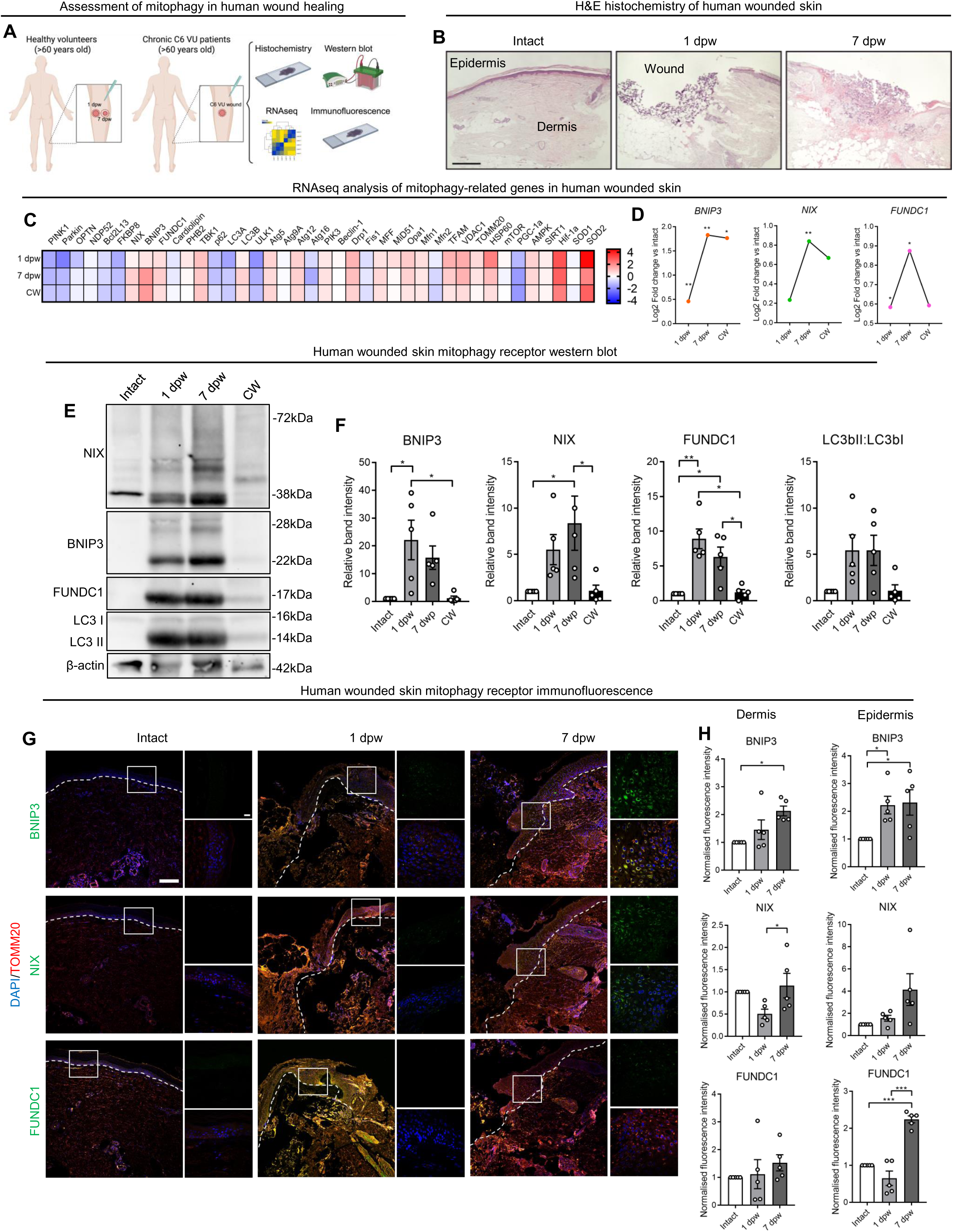
**Receptor-mediated mitophagy is upregulated in the early wound healing response.** (**A**) Schematic representation of wounded human skin biopsy acquisition. (**B**) Representative images of H&E-stained sections from intact, 1 dpw, 7 dpw, and CW samples. Scale bar = 500μm. (**C**) RNAseq profiling of mRNA gene expression of mitophagy-related genes from intact, 1 dpw, and 7 dpw (n = 22), as well as from venous ulcer chronic wound (CW) patients (n = 18). Gene expression is plotted as the log2fold change. (**D**) Graph of log2fold change mRNA expression of *BNIP3, NIX,* and *FUNDC1* in 1 dpw, 7 dpw, and CW tissue compared to intact. ** = p < 0.01; * = p < 0.05. (**E**) Representative immunoblot of mitophagy protein expression in full-thickness wound biopsies from acute wounds from healthy volunteers (n = 6), as well CW patients (n = 6). (**F**) Quantification (mean ± SEM) of relative protein abundance of mitophagy-related proteins in intact, 1 dpw, 7 dpw, and CW tissue (n = 6). Two-way ANOVA, ** = p < 0.01; * = p < 0.05. Each dot represents a biological replicate. (**G-H**) Immunofluorescence staining and (**G**) quantification (mean ± SEM) of NIX, BNIP3, and FUNDC1 fluorescence intensity in intact, 1 dpw, 7 dpw (all n = 5) biopsies. Two-way ANOVA, ** = p < 0.01; * = p < 0.05. Each dot represents a biological replicate.

Next western blot (WB) analysis of key receptor-mediated mitophagy markers in wounded skin lysates confirmed the significant upregulation of BNIP3 at 1 dpw, as well as BNIP3, NIX, and LC3-II at 7 dpw compared to intact skin (**Figures 1E and 1F**). Finally, immunofluorescence (IF) analysis of BNIP3, NIX, and FUNDC1 was performed on intact, 1 dpw, and 7 dpw skin sections (**Figure 1G**). Here, the expression of BNIP3 was significantly upregulated in both the dermis and epidermis at 7 dpw compared to intact skin, as well as at 1 dpw vs intact in the epidermis (**Figures 1G and 1H**). Additionally, the expression of FUNDC1 was significantly higher at 7 dpw compared to intact in the epidermis (**Figure 1H**). Collectively, these results demonstrated an upregulation of ub-independant mitophagy in the human skin wound healing response.

### Wounding induces elevated mitophagy, decreased mitochondrial mass, and mitochondrial fragmentation

As dermal fibroblasts perform important functions at the proliferation stage of wound healing, and mitophagy was primarily upregulated at this stage in human wounded skin tissue, we next investigated whether mitophagy was upregulated in primary human dermal fibroblasts (HDFb) *in vitro*. Here, following the creation of a wound gap between populations of HDFb, we performed live cell imaging of mitochondria and lysosomes in cells classified into three categories based on their spatial localisation to the wound gap – stationary; wound edge (WE); and migrating – at progressing time points (**Figures 2A and 2B**).

**Figure 2.**
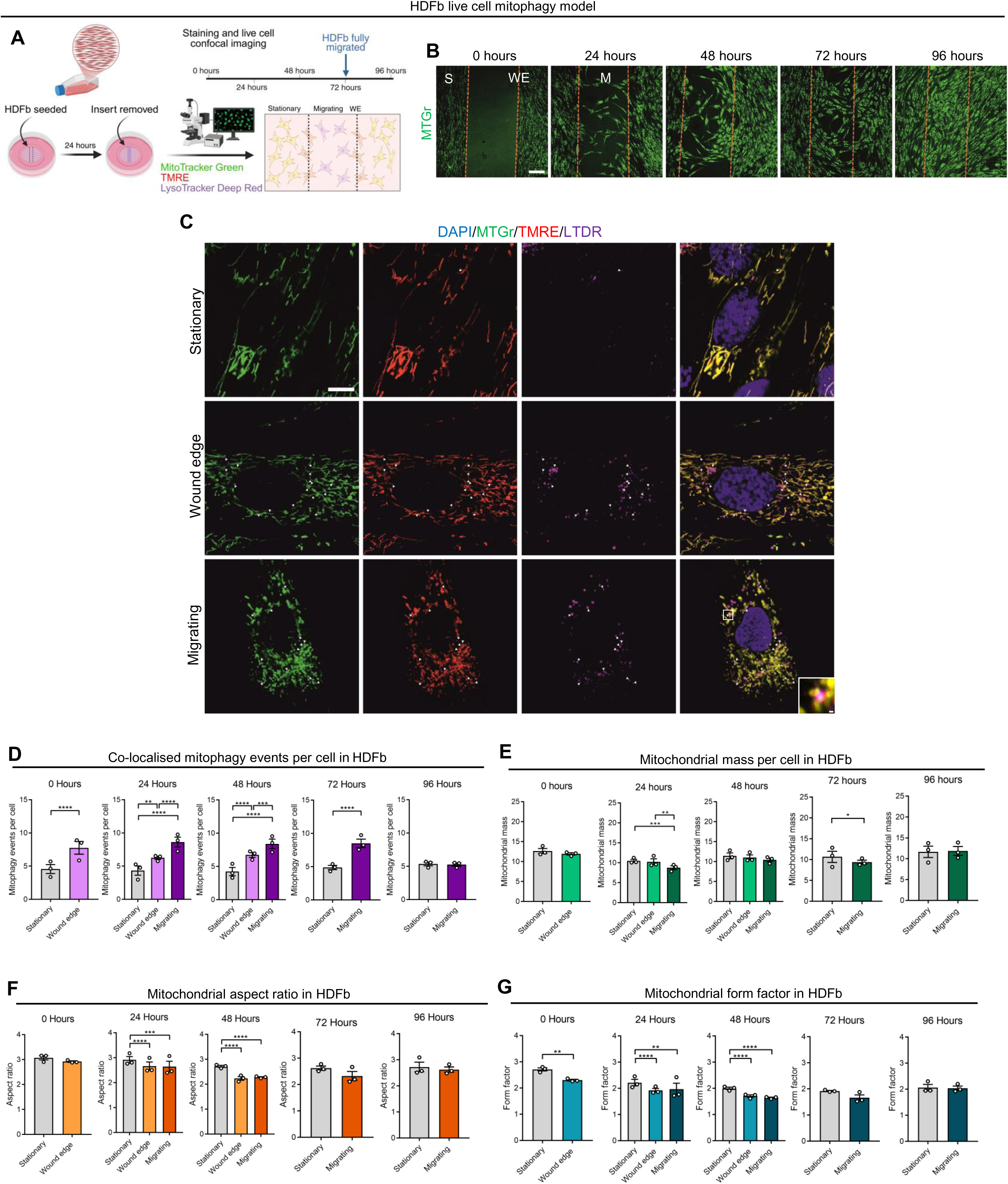
***In vitro* elevated mitophagy and mitochondrial fragmentation in migrating HDFb.** (**A**) Schematic representation of the experimental setup. (**B**) Representative fluorescence images of HDFb at 0, 24, 48, 72, and 96 hours in the live cell wounding assay. Orange lines represent the wound edge. Scale bar = 200µm. S = stationary; WE = wound edge; M = migratory. (**C**) Representative fluorescence images of intact, wound edge, and migrating HDFb stained with DAPI, MTGr, TMRE, and LTDR. Arrows depict colocalised lysosomes and mitochondria, representing mitophagy events. Scale bars = 10µm (full size), 0.5µm (zoom). (**D-G**) Quantification (mean ± SEM) of (**D**) co-localised mitophagy events, (**E**) mitochondrial mass, (**F**) mitochondrial aspect ratio, and (**G**) mitochondrial form factor in HDFb at sequential time points. Two-way ANOVA or students t-test, **** = p < 0.0001; *** = p < 0.001; ** = p < 0.01; * = p < 0.05. N = 30 cells in 3 separate biological replicates within each condition. Each dot represents the mean of an individual biological replicate.

When quantifying mitophagy events based on the co-localisation of lysosomes and mitochondria, it was found that WE cells had significantly higher levels of mitophagy events than intact cells at 0 hours, while at 24 hours and 48 hours, both WE and migrating cells had significantly higher levels of mitophagy compared to stationary cells. In addition, mitophagy was also significantly higher in migrating cells compared to WE cells at both 24 and 48 hours, and compared to stationary cells at 72 hours. There was no significant difference at 96 hours, when the cells had fully migrated (**Figures 2C and 2D**).

Next, when examining mitochondrial mass and morphology, mitochondrial mass was significantly reduced in migrating cells compared to both stationary and WE cells at 24 hours. Additionally, migrating HDFb had a significantly lower mitochondrial mass compared to stationary cells at 72 hours (**Figure 2E**). With regard to mitochondrial morphology, both aspect ratio (AR) and form factor (FF) were significantly decreased in both WE and migrating HDFb compared to stationary HDFb at the earlier time points prior to cessation of HDFb migration (**Figures 2F and 2G**). Taken together, these results suggest that both migrating and HDFb primed for migration localised at the WE have elevated mitophagy events, decreased mitochondrial mass, as well as contain more fragmented and less branched mitochondria.

### Urolithin A enhances fibroblast migration to accelerate wound healing

Following the demonstration that mitophagy is upregulated in the early and mid-stages of wound healing, we next sought to investigate whether pharmacologically enhancing mitophagy would accelerate wound healing. Firstly, we used human *ex vivo* full-thickness wound biopsies cultured for 7 days in various concentrations of three mitophagy inducers (UroA, nicotinamide riboside (NR), and BAM15) as well as the autophagy inhibitor chloroquine (CQ), and measured re-epithelialisation rate (**Figures 3A, S1A and S1B**). Here, re-epithelialisation rate was significantly increased in biopsies treated with all three concentrations of UroA, as well as 0.25mM NR compared to vehicle-treated biopsies. BAM15 had no effect on healing rate. Additionally, treatment with CQ at 50μM and 25μM significantly decreased healing rate (**Figures 3B, 3C, S1A, and S1B**). Specifically, healing rate was significantly enhanced in UroA-treated biopsies of different concentrations at 1, 3, and 5 dpw (**Figure S1C**).

**Figure 3.**
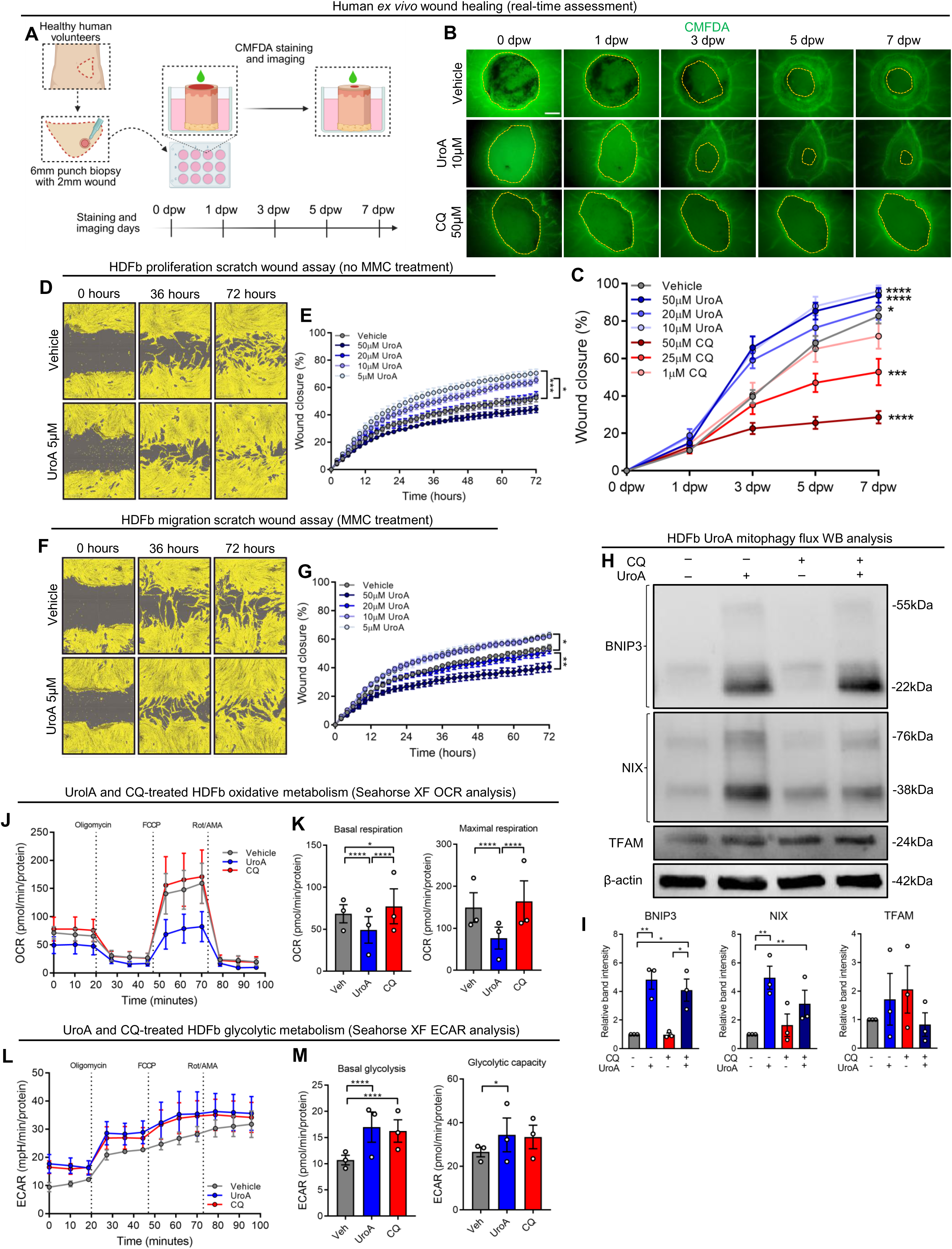
**Pharmacological induction of mitophagy increases HDFb migration.** (**A**) Schematic summary of *ex vivo* wound healing model. (**B**) Representative fluorescence images of wounded explant *ex vivo* biopsies treated with vehicle, 10μM UroA and 50μM CQ at 0, 1, 3, 5, and 7 dpw. Wound edge is indicated by the orange dotted line. Scale bar = 200μm. (**C**) Quantification (mean ± SEM) of wound healing time of wounds treated with various concentrations of UroA and CQ. Two-way ANOVA, **** = p < 0.0001; *** = p < 0.001, * = p < 0.05. N = 5 biological replicates. (**D-E**) Representative images and (**E**) quantification (mean ± SEM) of scratch wound assay in HDFb treated with vehicle and UroA. (**F-G**) Representative images and (**G**) quantification (mean ± SEM) of scratch wound assay in HDFb treated with vehicle + MMC and UroA + MMC over 72 hours. Two-way ANOVA, ** = p < 0.01, * = p < 0.05. N = 3 biological replicates containing at least 3 technical replicates. (**H**) Representative immunoblot analysis of BNIP3, NIX, and TFAM in HDFb. (**I**) Quantification (mean ± SEM) of BNIP3, NIX, and TFAM protein levels in HDFb. Two-way ANOVA, ** = p < 0.01, * = p < 0.05. N = 3 biological replicates. Each dot represents a biological replicate. (**J**) Quantification (mean ± SEM) of OCR assessment in vehicle, UroA, and CQ-treated HDFb. Dots represent the mean of 3 biological replicates, each containing at least 3 technical replicates. (**K**) Quantification (mean ± SEM) of basal respiration and maximal respiration in vehicle, UroA, and CQ-treated HDFb. Two-way ANOVA, **** = p < 0.0001, * = p < 0.05. N = 3 biological replicates. Each dot represents a biological replicate. (**L**) Quantification (mean ± SEM) of ECAR assessment in vehicle, UroA, and CQ-treated HDFb. Dots represent the mean of 3 biological replicates, each containing at least 3 technical replicates. (**M**) Quantification (mean ± SEM) of basal glycolysis and glycolytic capacity in vehicle, UroA, and CQ-treated HDFb. Two-way ANOVA, **** = p < 0.0001, * = p < 0.05. N = 3 biological replicates. Each dot represents a biological replicate.

As the effect of UroA was highest in *ex vivo* re-epithelialisation, we next investigated the effect of UroA on healing rate *in vitro* by performing scratch wound assays with HDFb treated with various concentrations of UroA and CQ. A subset of HDFb were additionally treated with mitomycin C (MMC) in order to inhibit proliferation and allow for the specific investigation of cell migration. Here, in MMC^-^ HDFb, UroA at 5μM and 10μM significantly accelerated wound closure compared to vehicle (**Figures 3D and 3E**), whilst 5μM UroA significantly increased healing rate in MMC^+^ HDFb compared to vehicle (**Figures 3F and 3G**). Of note, treatment with 50μM UroA significantly decreased healing rate in MMC^+^ HDFb compared to vehicle. Treatment with 50μM CQ also significantly decreased healing rate in MMC^-^ HDFb compared to vehicle (**Figure S1D and S1E**), as did both 50μM and 25μM in MMC^+^ HDFb (**Figure S1F and S1G**).

Next, in order to confirm the effect of UroA on receptor-mediated mitophagy in HDFb, WB analysis of BNIP3, NIX, and TFAM was undertaken in mitophagy flux settings whereby HDFb were additionally treated with 50μM CQ for 1 hour in pulse-chase setting, which demonstrated a significant upregulation of BNIP3 and NIX in UroA-treated HDFb, with no significant change in TFAM (**Figures 3H and 3I**). This was confirmed by live cell imaging of mitophagy events (**Figure S2A and S2B**), whilst caspase 3/7 (**Figure S2C and S1D**) and cell viability (**Figure S2E**) and assessment confirmed that both UroA and CQ had no long-term toxic effects on HDFb.

Finally, as previous studies^7^ have demonstrated the importance of a metabolic switch towards glycolysis in the early-stages of wound healing, we next investigated the impact of UroA on HDFb metabolism. Here, measurement of oxygen consumption rate (OCR) demonstrated a significant reduction in both basal and maximal respiration in UroA-treated HDFb (**Figures 3J and 3K**), whilst extracellular acidification rate (ECAR) measurement demonstrated a significant upregulation of basal glycolysis and glycolysis rate in UroA-treated HDFb (**Figures 3L and 3M**). Collectively, these results indicate that the mitophagy inducer UroA significantly increased migration of fibroblasts and accelerated wound healing, as well as inducing ub-independant mitophagy and switches to glycolytic metabolism.

### UroA enhances wound healing and collagen deposition in aged mouse model

In order to validate our findings that UroA increased wound healing in both *ex vivo* human explant biopsies as well as HDFb, we next examined the effect of UroA on wound healing *in vivo*. Here, full-thickness 6mm wound biopsies were taken from aged (>70 weeks) C57BL/6 mice treated orally with either 25mg/kg UroA or vehicle through gavage daily for 15 days.

There was no significant difference in body weight between vehicle and UroA-treated mice (**Figure S3A**). Healing rate was monitored every second day (**Figure 4A**), and macroscopic assessment of wound images demonstrated a significant acceleration in both wound healing and re-epithelialisation rate in UroA-treated mice compared to vehicle (**Figures 4B-4D**).

**Figure 4.**
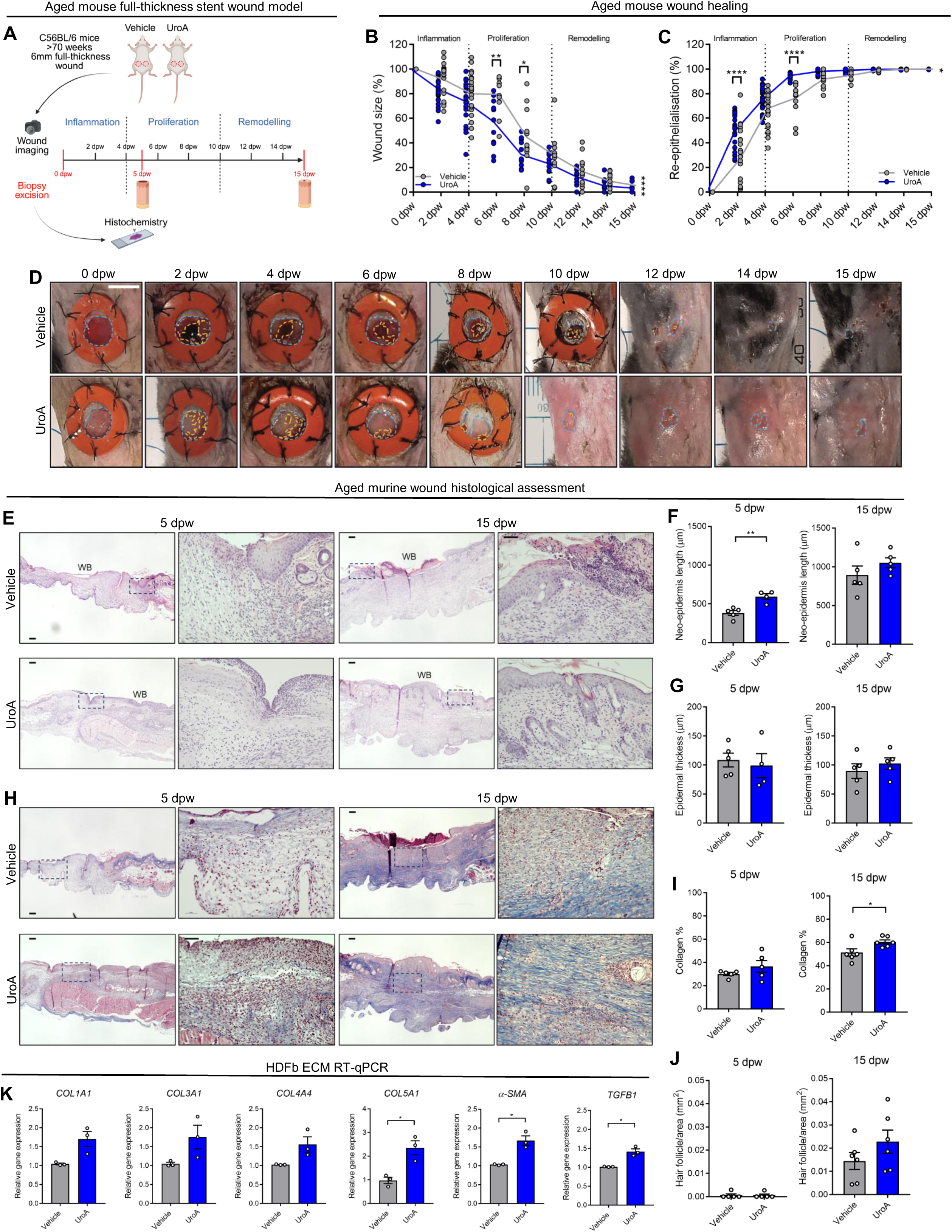
UroA enhances wound healing and collagen deposition in aged mouse model. (**A**) Schematic summary of aged mouse wound healing model. (**B-C**) Quantification (mean ± SEM) of (**B**) wound size (%) and (**C**) re-epithelialisation (%) in aged mice treated daily with either vehicle or UroA. Two-way ANOVA, **** = p < 0.0001, ** = p < 0.01, * = p < 0.05. Each dot represents an individual mouse (n = 24). (**D**) Representative photography images of wound healing in vehicle and UroA treated mice. Scale bar = 200μm. Blue line represents the wound area and orange line represents re-epithelialised area. (**E**) Representative images of H&E stained FFPE sections from vehicle and UroA-treated wounds at 5 dpw and 15 dpw. Scale bar = 200μm. WB = wound bed. (**F-G**) Quantification (mean ± SEM) of (**F**) neo-epidermis length (μm) and epidermal thickness (μm). Student’s t-test, ** = p < 0.01. Each dot represents an individual mouse (D5, n = 9; D15, n = 10). (**H**) Representative images of Masson’s trichrome stained FFPE sections from vehicle and UroA-treated wounds at 5 dpw and 15 dpw. Scale bar = 200μm. (**I-J**) Quantification (mean ± SEM) of (**I**) collagen (%) and (**J**) hair follicle abundance (per mm^2^). Student’s t-test, * = p < 0.05. Each dot represents an individual mouse (n = 24). (**K**) Quantification (mean ± SEM) of ECM-related genes in HDFb. Student’s t-test, * = p < 0.05. Each dot represents an individual biological repeats (n = 3).

Next, histological assessment of formalin-fixed paraffin embedded (FFPE) 5 and 15 dpw biopsies following H&E histochemical staining demonstrated a significant increase in neo-epidermis length in UroA-treated 5 dpw biopsies when compared to vehicle (**Figures 4E and 4F**), whilst there was no significant difference in epidermal thickness (**Figure 4G**).

Additionally, Masson’s trichome histology showed an increase in collagen deposition in the neo-dermis within 15 dpw in UroA-treated biopsies when compared to vehicle (**Figures 4H and 4I**). There was no significant difference in the abundance of newly-formed hair follicles in the neo-dermis (**Figure 4J**). As a proof of principle that UroA induces collagen synthesis, we then performed RT-qPCR analysis of various ECM-related genes in HDFb. Here, UroA treatment significantly increased the expression of *COL5A1, α-SMA,* and *TGFB1* in HDFb compared to vehicle (**Figure 4K**), whilst there was no significant difference in the expression of *COL1A1, COL3A1, COL4A4* (**Figure 4K**), *TGFA, MMP2, MMP9,* and *MMP10* (**Figure S3B**).

### UroA upregulates non-canonical WNT5A signalling to accelerate fibroblast migration

To investigate the cellular and molecular mechanisms by which UroA upregulated migration in HDFb, we next undertook RNAseq analysis of HDFb treated either with 5μM UroA or vehicle for 24 hours. PCA plots confirmed a variance based on treatment type (**Figure S4A**). Next, DEG and gene enrichment analysis revealed an upregulation in genes involved in non-canonical WNT signalling in UroA-treated HDFb, including an upregulation in ‘Ca^2+^ pathway’, ‘Signaling By WNT’, ‘WNT Ligand Biogenesis And Trafficking’, and ‘Beta-catenin Independent WNT Signaling’ (**Figures 5A and 5B**).

**Figure 5.**
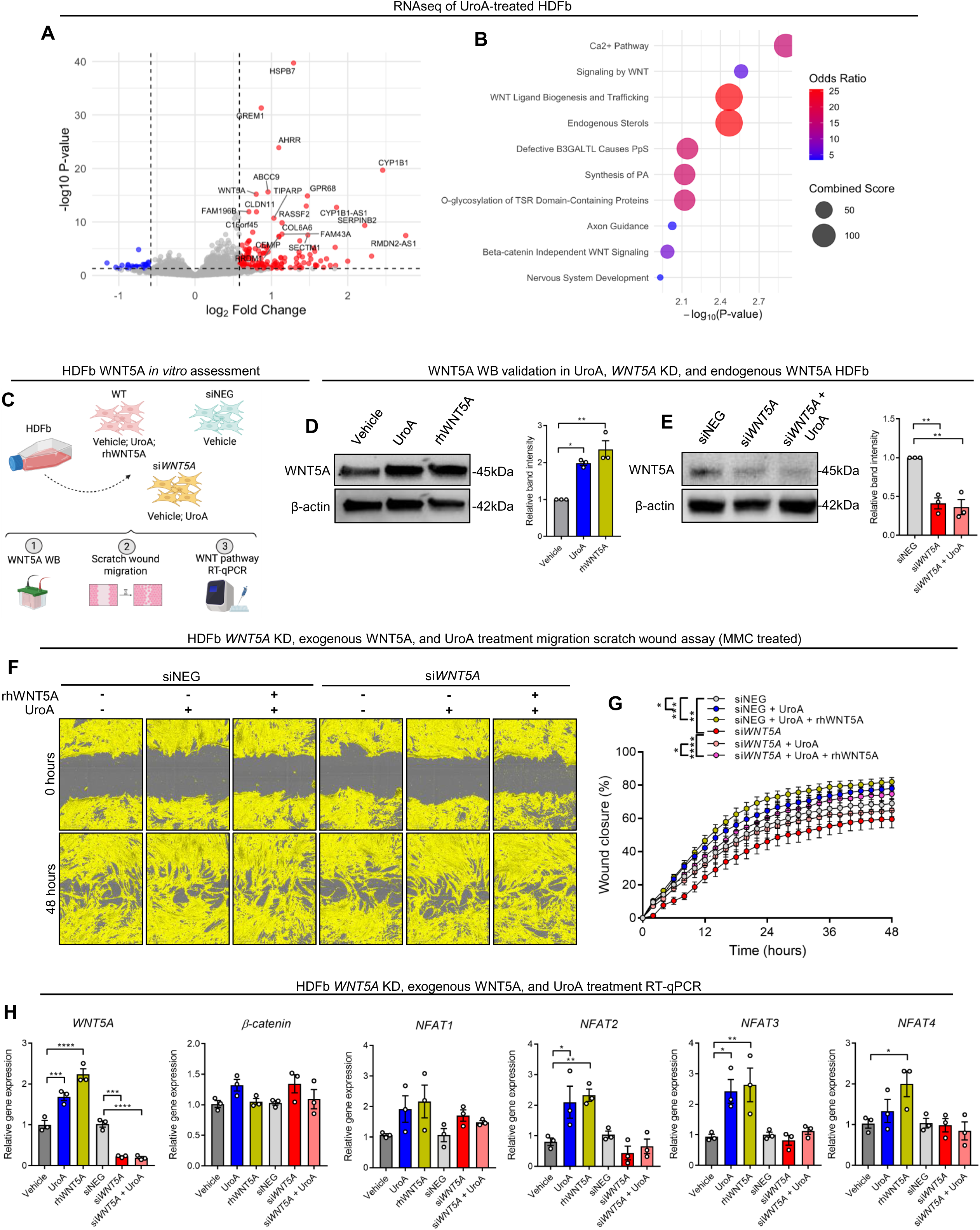
**UroA upregulates non-canonical WNT5A signalling to accelerate fibroblast migration** (**A**) Volcano plot depicting DEG quantification of UroA vs vehicle HDFb gene expression. Genes significantly upregulated in UroA HDFb (red) and significantly downregulated (blue) compared to vehicle HDFb. (**B**) Gene enrichment analysis showing the 10 top significantly upregulated pathways in UroA treated HDFb from Reactome database. (**C**) Schematic summary of the *in vitro* experimental assessment. (**D-E**) Representative immunoblot and quantification (mean ± SEM) of relative WNT5A protein levels of WNT5A in (**D**) WT and (**E**) siNEG and si*WNT5A* HDFb. Two-way ANOVA, ** = p < 0.01, * = p < 0.05. Each dot represents an individual biological replicate (n = 3). (**F**) Representative phase contrast images of HDFb at 0 hour and 48 hours post scratching in scratch wound migration assessment. (**G**) Quantification (mean ± SEM) of HDFb migration in scratch wound migration analysis. Two-way ANOVA, **** = p < 0.0001. Each dot represents the mean of 3 biological replicate which each contain at least 2 technical replicates. (**H**) Quantification (mean ± SEM) of RT-qPCR analysis of WNT pathway genes in HDFb. Two-way ANOVA, **** = p < 0.0001, *** = p < 0.001, ** = p < 0.01, * = p < 0.05. Each dot represents an individual biological replicate.

We next sought to further elaborate on the association between UroA and WNT5A in HDFb (**Figure 5C**). Firstly, WB analysis confirmed the significant increase in WNT5A protein levels in UroA-treated HDFb, as well HDFb treated exogenously with recombinant human WNT5A (rhWNT5A) for 24 hours (**Figure 5D**), whilst WNT5A was significantly decreased in HDFb with transient *WNT5A* knockdown (KD, si*WNT5A*) compared to HDFb transfected with negative control (siNEG) (**Figure 5E**). To determine the importance of WNT5A in HDFb migration, as well as whether treatment with exogenous WNT5A could rescue the UroA-induced migration abolished by *WNT5A* KD, scratch wound assay was undertaken in HDFb pre-treated with MMC. Here, both UroA and rhWNT5A treated siNEG HDFb had a significantly elevated healing rate compared to vehicle. Importantly, whilst si*WNT5A* HDFb had a significantly decreased healing rate when compared to siNEG, treatment of si*WNT5A* with UroA failed to increase HDFb migration, although additional treatment with rhWNT5A rescued migration in si*WNT5A* (**Figures 5F and 5G**) – thereby demonstrating that Uro-A effect on mitophagy is mediated through WNT5A. Supporting the idea that WNT5A is important for fibroblast function during the proliferation stage of wound healing, Examination of previously-published single-cell RNAseq (scRNAseq) data^23^ revealed the increased expression of both *WNT5A* and its long non-coding RNA, *WNT5A-AS1*, in fibroblast subsets at day 7 post wounding tissue (**Figure S4B**).

As there are various known arms of WNT pathways involving WNT5A, both canonical and non-canonical, we next assessed the gene expression of various genes associated with either canonical WNT5A, non-canonical WNT5A/planar cell polarity (PCP), or non-canonical WNT5A/Ca^2+^ pathways. Here RT-qPCR analysis demonstrated the significant upregulation in *NFAT2* and *NFAT3* expression in UroA-treated HDFb compared to vehicle, as well as the expression of *NFAT2, NFAT3,* and *NFAT4* in rhWNT5A-treated HDFb compared to vehicle (**Figure 5H**). As there was no significant difference in the gene expression in any of the genes associated with either canonical WNT5A or non-canonical WNT5A/PCP pathways (**Figure S4C**), these results collectively suggested that UroA was accelerating HDFb migration through the WNT5A/Ca^2+^ pathway.

### UroA-induced mitophagy upregulates CREB-mediated transcription of *WNT5A* through cytosolic Ca^2+^ elevation

Mitochondria are known stores of intracellular Ca^2+^, and the degradation of mitochondria through mitophagy has previously been shown to increase cytosolic Ca^2+^ levels. To test this hypothesis, we firstly confirmed the importance of Ca^2+^ in HDFb migration through scratch wound analysis of MMC^+^ HDFb treated with various concentrations of the cytosolic Ca^2+^ quencher BAPTA-AM, which demonstrated a significant decrease in wound closure compared to vehicle (**Figures 6A and 6B**). Cell viability assessment showed no significance difference in cell viability, confirming that the decrease in migration was not due to any potential toxic effects of BAPTA-AM (**Figure S5A**).

**Figure 6.**
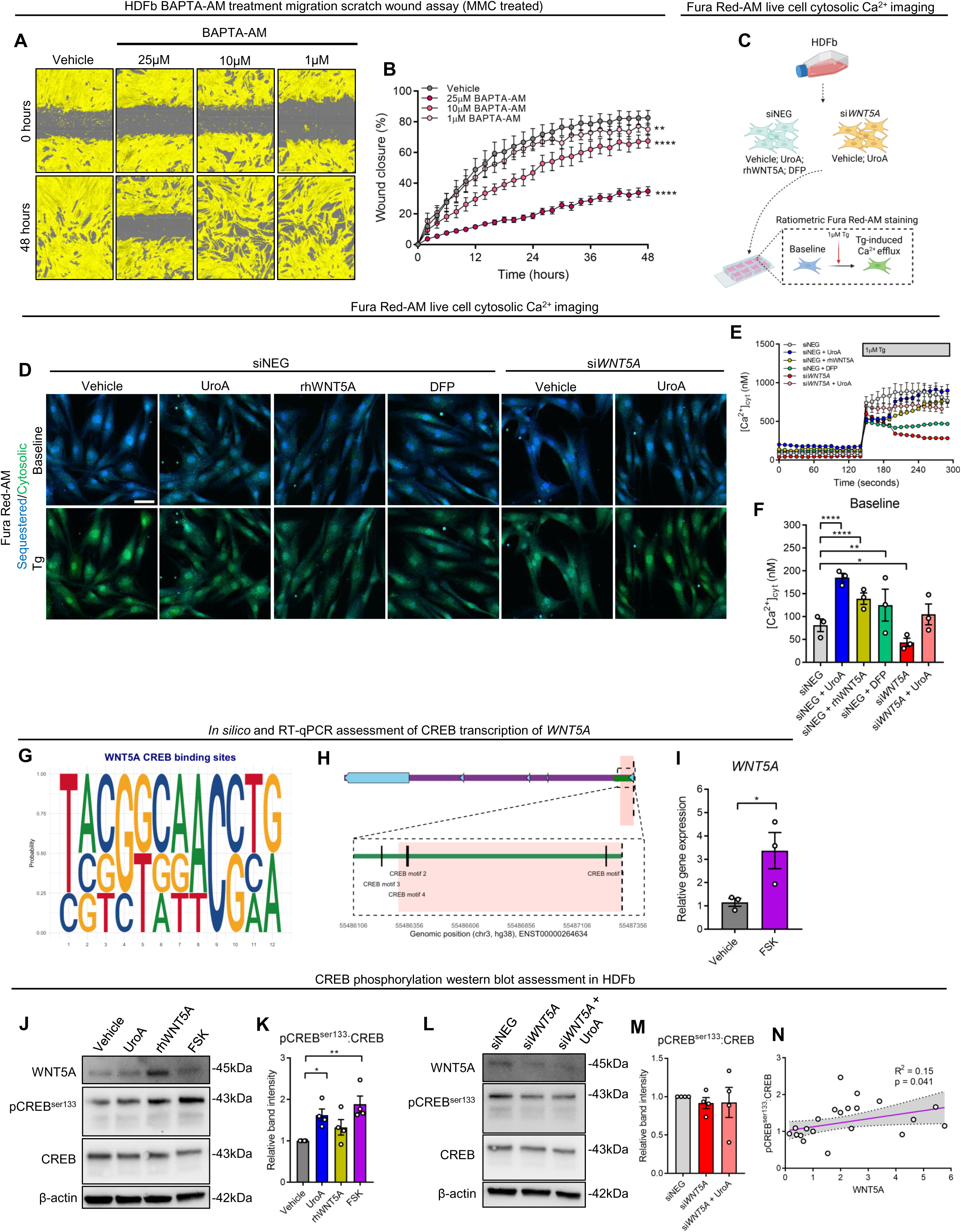
**UroA-induced mitophagy upregulates CREB-mediated transcription of *WNT5A* through cytosolic Ca^2+^ elevation** (**A**) Representative phase contrast images of HDFb incubated with either vehicle or BAPTA-AM at 0 hour and 48 hours post scratching in scratch wound migration assessment. (**B**) Quantification (mean ± SEM) of HDFb migration in scratch wound migration analysis. Two-way ANOVA, **** = p < 0.0001, ** = p < 0.01. Each dot represents the mean of 3 biological replicate with at least 2 technical replicates. (**C**) Schematic summary of live cell cytosolic Ca^2+^ imaging in HDFb. (**D**) Representative fluorescence images of HDFb stained with Fura Red-AM at baseline and following 1μM Tg treatment. Scale bar = 10µm. (**E**) Quantification (mean ± SEM) of [Ca^2+^]_cyt_ (nM) in HDFb. Each dot represents the mean of 3 biological replicates containing at least 20 cells each. (**F**) Quantification (mean ± SEM) of baseline [Ca^2+^]_cyt_ (nM) in HDFb. Two-way ANOVA, **** = p < 0.0001, ** = p < 0.01, * = p < 0.05. (**G**) Motif analysis of putative CREB binding sites in the *WNT5A* promoter region. (**H**) Genomics viewer of *WNT5A* gene (purple) with the transcription starter sequence (red), exons (cyan), promoter region (green), and putative CREB binding sites. (**I**) Quantification (mean ± SEM) of *WNT5A* gene expression in vehicle and FSK treated HDFb. Student’s t-test, * = p < 0.05. Each dot represents an individual biological replicate. (**J-M**) Quantification (mean ± SEM) and (**K, M**) representative immunoblot of pCREB^ser1^^33^, CREB, and WNT5A in HDFb. Two-way ANOVA, ** = p < 0.01, * = p < 0.05. Each dot represents an individual biological replicate. (**N**) Linear regression analyses of the correlation between pCREB^ser1^^33^/CREB ratio and WNT5A protein levels. Each dot represents an individual replicate (N = 3 biological replicates per condition.

Next, to investigate whether enhanced mitophagy through UroA treatment led to an increase in cytosolic Ca^2+^ levels, we performed live cell imaging of HDFb transfected with either siNEG or si*WNT5A* following staining with the ratiometric cytosolic Ca^2+^ indicator Fura Red-AM (**Figure 6D**). Optimisation of *Rmin* (50μM BAPTA-AM) and *Rmax* (1μM ionomycin) allowed for the quantification of absolute cyctosolic Ca^2+^ (**Figure S5B and S5C**). Here, baseline cytosolic Ca^2+^ levels were significantly higher in siNEG HDFb pre-treated with either UroA, rhWNT5A, or the iron chelator deferiprone (DFP) – a potent mitophagy inducer (**Figures 6D-F**). In addition, baseline cytosolic Ca^2+^ was significantly lower in si*WNT5A* compared to siNEG HDFb. 1μM thapsigardin (Tg) was then added to induce Ca^2+^ efflux and allow for the measurement of Ca^2+^ buffering capacity. Although Tg-induced Ca^2+^ efflux was significantly decreased in DFP-treated siNEG HDFb compared to siNEG, there was no significant difference in either UroA- or rhWNT5A-treated siNEG HDFb compared to siNEG (**Figure S5D**), suggesting that although baseline levels of cytosolic Ca^2+^ are higher in these conditions, there is no impact on mitochondrial Ca^2+^ stores.

We next sought to investigate how increases in cytosolic Ca^2+^ levels may lead to enhanced *WNT5A* transcription. RT-qPCR analysis of *WNT5A* and *NFAT* mRNA levels demonstrated a significant increase in *WNT5A*, *NFAT1, NFAT2,* and *NFAT3* in HDFb treated with 1mM ionomycin (**Figure S5E**). Next, as previously both ionomycin-induced cytosolic Ca^2+^ elevation and UroA treatment significantly increased *NFAT* gene expression in HDFb, we hypothesised that UroA upregulated *WNT5A* transcription through catalysing the de-phosphorylation and nuclear translocation of NFAT3. However, WB quantification showed no significant increase in NFAT3 de-phosphorylation in UroA-treated HDFb compared to vehicle (**Figure S5F**).

Subsequently, we next investigated whether activation of the Ca^2+^-sensitive cAMP Response Element-Binding Protein (CREB) led to increased *WNT5A* transcription. Firstly, through *in silico* analysis using JASPAR motif analysis, we observed the presence of four separate putative CREB-binding sites with relative scores exceeding 80% located within or near the *WNT5A* promoter sequence (**Figures 6G, 6H and S5G**). RT-qPCR analysis demonstrated that treatment of HDFb with 10μM forskolin (FSK) – a direct activator of adenylyl cyclase and CREB phosphorylation – significantly increased *WNT5A* mRNA levels compared to vehicle (**Figure 6I**). Next, WB quantification confirmed that treatment of HDFb with both UroA and rhWNT5A significantly increased CREB phosphorylation, as measured by the ratio of CREB:pCREB^ser133^, when compared to vehicle (**Figures 6J and 6K**). KD of *WNT5A* did not decrease pCREBser1313 levels (**Figure 6L and 6M**). Importantly, treatment with FSK significantly increased WNT5A as well as pCREB^ser133^ protein levels compared to vehicle, and linear regression analysis confirmed a statistically significant positive correlation between pCREB^ser133^ and WNT5A (**Figure 6N**). Altogether, these results collectively suggest that elevated cytosolic Ca^2+^ levels induced by UroA-mediated mitophagy led to an upregulation of *WNT5A* transcription through increased CREB phosphorylation.

### UroA-mediated WNT5A/Ca^2+^ signalling upregulates actin polymerisation to enhance fibroblast migration

WNT5A signalling has previously been shown to promote cellular migration through enhancing actin polymerisation. In order to determine whether UroA-mediated WNT5A/Ca^2+^ signalling was important for actin polymerisation in HDFb, we firstly performed immunofluorescence (IF) staining for F-actin capping protein. Here, both UroA- and rhWNT5A-treated HDFb exhibited significantly decreased levels of F-actin capping protein compared to vehicle, whilst si*WNT5A* and si*WNT5A* treated with UroA HDFb displayed significantly increased F-actin capping protein levels compared to siNEG (**Figures 7A and 7B**). As F-actin capping protein is known to indicate non-active actin polymerisation, these results suggest that WNT5A was important for actin polymerisation.

**Figure 7.**
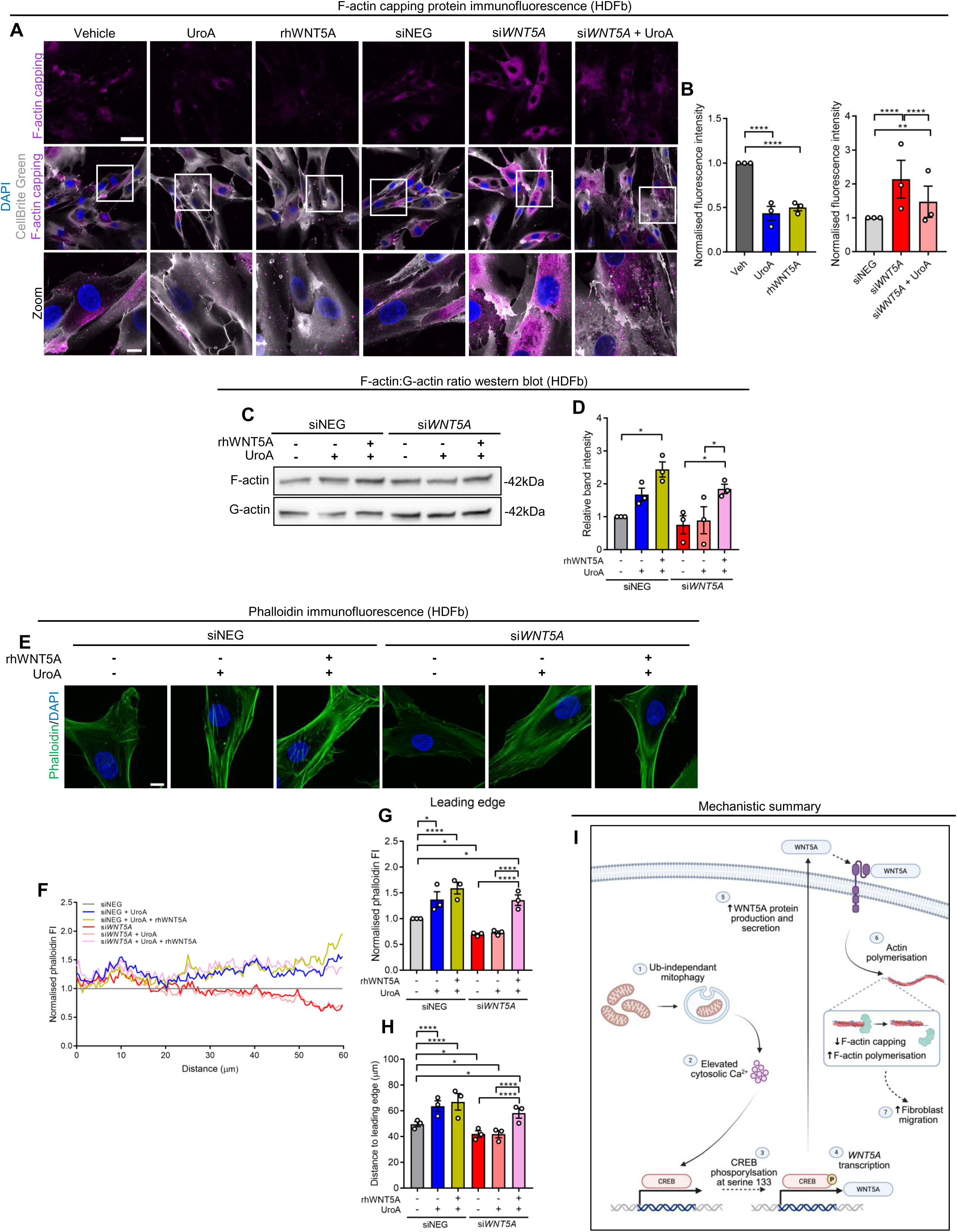
**UroA-mediated WNT5A/Ca^2+^ signalling upregulates actin polymerisation to enhance fibroblast migration** (**A**) Representative immunofluorescence images of F-actin capping protein in HDFb. Scale bar = 50μm (full size), 10μm (zoom). (**B**) Quantification (mean ± SEM) of F-actin capping protein fluorescence intensity. Two-way ANOVA, **** = p < 0.0001, * = p < 0.05. Each dot represents an individual biological replicate containing at least 15 cells. (**C**) Representative immunoblot of F-actin and G-actin in HDFb. (**D**) Quantification (mean ± SEM) of F-actin:G-actin ratio in HDFb. Two-way ANOVA, ** = p < 0.01, * = p < 0.05. Each dot represents an individual biological replicate. (**E**) Representative immunofluorescence images of phalloidin staining in HDFb. Scale bar = 10µm. (**F**) Point plot of phalloidin fluorescence intensity normalised to siNEG (grey) in HDFb. Each line represents the mean of 3 biological replicated each containing 15 cells. 60μm indicates the cell periphery. (**G-H**) Quantification (mean ± SEM) of (**G**) the average phalloidin fluorescence intensity at the final 10μm of the cell periphery and (**H**) distance (μm) from the nuclei to the periphery of the leading edge. Two-way ANOVA, **** = p < 0.0001, * = p < 0.05. Each dot represents an individual biological replicate containing 15 individual cells. (**I**) Schematic summary of the mechanism by which UroA-mitophagy increases WNT5A/Ca^2+^ signalling.

To confirm this, we next performed WB to quantify the ratio of G-actin:F-actin. Here, rhWNT5A treatment significantly increased the F-actin ratio in siNEG (**Figures 7C and 7D**). Importantly, si*WNT5A* treated with both UroA and rhWNT5A displayed significantly elevated F-actin ratio levels compared to si*WNT5A*, whilst si*WNT5A* treated only with UroA did not rescue HDFb migration, suggesting that UroA acts upstream of *WNT5A* transcription. F-actin was higher in siNEG treated with UroA, although this was not significant, whilst F-actin ratio was significantly higher in wild-type HDFb treated with UroA, rhWNT5A, and recombinant Epidermal Growth Factor (EGF) (**Figures S6A and S6B**). Next, to assess the spatial localisation of F-actin in HDFb, we performed IF staining of phalloidin, a component specific to the active form of actin, F-actin (**Figures 7E and 7F**). Here, treatment of HDFb with either UroA, rhWNT5A, or EGF significantly increased the levels of F-actin at the leading edge (**Figures S6C and S6D**). Phalloidin intensity was significantly higher at the leading edge in both UroA- and rhWNT5A-treated siNEG compared to siNEG, and was significantly lower in si*WNT5A* compared to siNEG. Importantly, phalloidin IF was significantly higher in si*WNT5A* treated with both UroA and rhWNT5A compared to both si*WNT5A* and si*WNT5A* + UroA, again suggesting that UroA acts upstream of *WNT5A* transcription (**Figures 7G and 7H**).

Quantification of the distance of the leading edge to cell nucleus displayed similar trends as with phalloidin intensity (**Figures 7H and S6E and S6F**), supporting the idea that UroA increased HDFb motility through elevated actin polymerisation at the leading edge, and that UroA acts upstream of *WNT5A* transcription (**Figure 7I**).

## Discussion

In this study, we demonstrate that ub-independant mitophagy, particularly BNIP3- and NIX-mediated mitophagy, is upregulated in the inflammation and proliferation stages of human wound healing. By promoting mitophagy with the well-characterised inducer UroA, we showed that mitophagy induction promoted wound healing by accelerating fibroblast migration. This occurred through the upregulation of non-canonical WNT5A signalling in a Ca^2+^-sensitive signalling pathway manner mediated through CREB phosphorylation, leading to enhanced actin polymerisation and cell motility. These findings thus uncover a link between mitophagy and non-canonical WNT signalling, demonstrate for the first time an important role for mitophagy in fibroblast function during wound healing, and suggest the potential benefit of targeting mitochondrial-related functions therapeutically in the wound healing context.

Mitochondrial quality control mechanisms, including mitophagy, have been increasingly recognised as an important regulator of various mitochondrial and cellular functions, including cellular metabolism^24,25^, signalling pathway modulation^26^, and apoptosis^27^. Indeed, mitochondria play essential roles in normal wound healing function^14^. Although BNIP3- and NIX-mediated mitophagy have previously been shown to accelerate both the migration^17,18^ and differentiation^21^ of keratinocytes in a non-wound healing context, no studies have investigated the role of mitophagy in human samples of wound healing, or its role in fibroblast function during wound healing. As such, the demonstration that BNIP3- and NIX-mediated mitophagy are upregulated at the inflammation and proliferation stage of wound healing in human wounded tissue both adds to and reinforces the knowledge that mitochondria play important roles in various aspects of wound healing^14^.

UroA is a well characterised mitophagy inducer derived from multi-step metabolism of dietary ellagitannins in the human gut microbiota^28^. Unlike other pharmacological mitophagy inducers, treatment with UroA has been shown to be safe for humans^29^, and previous studies have demonstrated its ability to improve the function of tissues in various disease contexts^29,30^ such as skin ageing^31,32^, muscle in Duchenne muscular dystrophy^33,34^, cardiovascular health^32^, and improved angiogenesis in wound healing^16^. Indeed, using both human *ex vivo* explant and *in vivo* mouse models of wound healing, we demonstrated that lower concentrations of UroA treatment accelerated wound closure and re-epithelialisation. The discrepancy in sensitivity between full-thickness explant biopsies and *in vitro* monolayers most likely explains the fact that 50μM UroA decreased fibroblast migration here but not in the *ex vivo* experiments.

During the proliferation stage of wound healing, fibroblasts migrate to the provisional wound matrix and differentiate into myofibroblasts, depositing collagen and facilitating extracellular matrix remodelling^5^. *In vitro* scratch wound migration assessment of primary dermal fibroblasts confirmed the motility-promoting actions of UroA on dermal fibroblasts, whilst collagen deposition was increased in the ECM of UroA-treated mice, and the gene expression of *COL5A1, α-SMA,* and *TGFB1* was significantly upregulated in UroA-treated HDFb. Previous studies have shown that metabolic programming towards a glycolytic phenotype accelerated the migration of fibroblasts during wound healing^7^, whilst the mitochondrial membrane protein cyclophilin D contributed to fibroblast-mediated collagen deposition during wound healing^35^. As such, our data demonstrating an upregulation in glycolytic metabolism and collagen deposition suggested an important link for mitophagy in these contexts.

Gene expression analysis of UroA-treated HDFb demonstrated an upregulation in non-canonical WNT5A signalling, which was confirmed at the protein level. WNT5A is one of 19 members of the WNT family of proteins, and is predominantly involved in activating the β-catenin-independent signalling cascades^36^. These non-canonical signalling pathways are known to play important roles in wound healing by promoting stem cell activity^37^, hair follicle morphogenesis^38^, and extracellular remodelling^39^, as well as in promoting fibroblast migration, differentiation, and TGF-β signalling in non-wound healing contexts^40^. Non-canonical WNT5A signalling can broadly be divided into the PCP and Ca^2+^ pathways, in which there are various modulators and effectors involved^41,42^. Subsequent RT-qPCR analysis of various known effectors suggested the activation of the Ca^2+^ pathway – as demonstrated by the upregulation in various *NFAT* mRNA levels – whilst scratch wound migration confirmed the migratory-enhancing capabilities of WNT5A signalling in HDFb.

Ca^2+^ signalling is an essential mediator of numerous cell signalling pathways^43^, including gene expression, metabolism, differentiation, and cell death pathways^44^, as well as in activating N-cadherin to promote fibroblast migration and cell-cell adhesion during wound healing^45^, whilst mitochondria are important Ca^2+^ sinks responsible for mediating Ca^2+^ buffering^46^. Indeed, the interplay between Ca^2+^ homeostasis and mitophagy is well described^43,47^, and in response to stressors, the degradation of mitochondria through mitophagy stimulates the elevation of cytosolic Ca^2+^ levels, as well as promoting Ca^2+^ release from endoplasmic reticulum (ER) following the breakdown of mitochondria-ER contact sites (MERCs)^48,49^. Our data demonstrated firstly that quenching of cytosolic Ca^2+^ inhibited the migration of primary dermal fibroblasts. Through live cell imaging of cytosolic Ca^2+^, we also showed that the induction of mitophagy through UroA and DFP treatment led to rises in baseline cytosolic Ca^2+^ levels. Interestingly, whilst DFP treatment impeded Tg-induced Ca^2+^ release, UroA treatment did not, suggesting that UroA-induced mitophagy did not significantly deplete Ca^2+^ stores. The fact that RT-qPCR analysis indicated that *WNT5A* transcription was upregulated in response to ionomycin-induced cytosolic Ca^2+^ elevation reinforced the theory that *WNT5A* transcription was mediated in a Ca^2+^-dependant signalling manner. However, WB analysis demonstrated no increase in NFAT3 de-phosphorylation.

CREB is a Ca^2+^-dependant transcription factor known to promote the transcription of various genes involved in cell proliferation, differentiation, survival, and motility^50^. This occurs through the binding of phosphorylated CREB to cAMP response elements (CRE) within their respective promoter regions^51^, and has been shown to be important in macrophage function both during wound healing^52^ and neuronal development^53^. However, whilst previous studies have demonstrated a non-direct link between CREB and non-canonical WNT signalling pathways^54^, no such studies have linked CREB directly to *WNT5A* transcription. As CREB is commonly activated by Ca^2+^-sensitive kinases such as protein kinase A (PKA), protein kinase C (PKC), and calcium/calmodulin-dependant protein kinase II (CaMKII)^55,56^, we hypothesised that cytosolic Ca^2+^ elevation in response to UroA-induced mitophagy may lead to *WNT5A* transcription through CREB activity. Accordingly, we first undertook *in silico* analysis of the *WNT5A* promoter region, which determined the existence of four putative CREB binding sites – suggesting the existence of functional CREs here. At the protein level, UroA treatment significantly increased CREB phosphorylation at serine 133, and there was a statistically significant positive correlation between pCREB and WNT5A protein levels, whilst treatment with the known CREB activator FSK also led to elevated *WNT5A* mRNA levels.

Collectively, these results suggested that UroA treatment induced the transcriptional upregulation of *WNT5A* by promoting the release of cytosolic Ca^2+^, followed by the subsequent phosphorylation of CREB and binding of CREs to the *WNT5A* promoter region. Accordingly, these results provide evidence of a direct link between Ca^2+^-mediated CREB pathway signalling and *WNT5A* transcription, as well as establishing a connection between mitophagy and non-canonical WNT5A signalling.

After demonstrating how UroA-induced mitophagy led to the transcriptional upregulation of *WNT5A*, we next sought to determine how elevated WNT5A promoted the migration of dermal fibroblasts. For motility, fibroblasts depend on an intricate, rapid, and tightly choregraphed process of cycles of actin polymerisation at the barbed end and depolymerisation at the pointed end^57^, resulting in the growth of filopodia and lamellipodia at the barbed end and subsequently migration^58,59^. Previous studies have demonstrated a role of WNT5A in inducing cytoskeletal rearrangements – in the form of concurrent increases in F-actin and decreases in vimentin and microtubule formation – in fibroblasts within fibrotic disease models^40^. Through immunofluorescence staining of F-actin capping protein, a protein which binds to the barbed end of F-actin tubules to stall polymerisation^60^, we demonstrated that treatment with both UroA and rhWNT5A significantly decreased F-actin capping protein abundance, whilst KD of *WNT5A* ultimately increased levels – indicating the enhancing effect of both UroA and WNT5A on F-actin polymerisation. Next, quantification of the abundance of the active, F-actin, form of actin through WB analysis of F-actin to G-actin ratio demonstrated that WNT5A increased F-actin levels in both siNEG and si*WNT5A* HDFb, whilst IF of phalloidin confirmed the increased expression of F-actin at the leading edge of HDFb treated with UroA and rhWNT5A. Interestingly, in both the WB and IF assessment of F-actin, the inhibiting effects of *WNT5A* KD was not rescued by UroA treatment, although was by the additional treatment with exogenous WNT5A protein. This suggested that UroA acted upstream of WNT5A protein expression and primarily on *WNT5A* transcription. Confirming this, scratch wound migration analysis demonstrated that UroA treatment of si*WNT5A* did not prevent the abolishment of migration seen with *WNT5A* KD, although additional treatment with exogenous WNT5A did so, indicating that UroA acts upstream of *WNT5A* transcription, endogenous protein production, and WNT5A-mediated migration.

In summary, our findings identify a previously unrecognised mechanism by which mitophagy, and in particular UroA-upregulated ub-independant mitophagy, promotes fibroblast migration via the non-canonical WNT5A/Ca^2+^ signalling pathway, offering new insights into the roles of mitochondria in wound healing, as well as the intersection of mitochondrial quality control and fibroblast function during wound healing. These findings may have therapeutic implications. As fibroblast migration is a key event in dermal wound closure, and impairment of this function contributes to the formation of chronic wounds, UroA could be a promising therapeutic candidate for enhancing wound repair. In addition, our findings are the first to directly show that cytosolic Ca^2+^ increases directly lead to *WNT5A* transcription, as opposed to simply being part of a feedback mechanism. Thus, the WNT5A/Ca^2+^ pathway may also serve as a mechanistic target for modulating fibroblast behaviour in both physiological and pathological contexts.

## Methods

### Sex as a biological variable

Our study examined male and female human wound healing and female mice. Sex was not considered a biological variable.

### Ethics

This study was approved by Stockholm Regional Ethics Committee and executed in agreement with the Helsinki Declaration. Informed consent was obtained from all the research subjects.

### Human samples

Healthy volunteers (>60 years of age) were enrolled at Karolinska University Hospital Dermatology Clinic, Stockholm, Sweden. Full-thickness skin wounds were made with a 4-mm biopsy punch in two adjacent spots 2 cm apart on the distal lower leg of the healthy volunteers. On 1 and 7 dpw, the wound edge area was excised with a 6-mm biopsy punch from one of the existing wounds. Similar wound edge biopsies were obtained on one occasion from patients (>60 years of age) with chronic C6 venous ulcers for >3 months^61^. All samples were either snap frozen in liquid nitrogen or fixed in 4% formalin (FFPE) and used for downstream analysis.

### Cell culture and treatments

HDFb were isolated from breast tissue of healthy adults and cultured in DMEM glucose free media (Thermo Fisher Scientific) supplemented with 10% Fetal Bovine Serum (FBS) (Thermo Fisher Scientific), 10mM glucose and antibiotics (100 units/mL Penicillin and 100 µg/mL Streptomycin (Thermo Fisher Scientific)) in an incubator maintained at 37℃ and 5% CO_2_. HDFb were treated with a range of concentrations of UroA and CQ (as stated in the more detail in the relevant section), as well as 1mM DFP (Santa Cruz Biotechnology), and a range of concentrations of both NR (Sigma-Aldrich) and BAM-15 (Sigma-Aldrich) for 24 hours. HDFb were treated with 250ng/ml rhWNT5A (R&D Systems) for 24 hours as a positive control for WNT5A. In *WNT5A* transcription experiments, HDFb were treated with various concentration of BAPTA-AM (R&D Systems) for up to 48 hours in order to quench cytosolic Ca^2+^ levels; 100nM FK506 (Invitrogen) for 24 hours to inhibit NFAT3 de-phosphorylation; as well as 10μM FSK (Merck) for 1 hour as a positive control for CREB phosphorylation. For actin polymerization experiments, HDFb were treated with 100ng/ml recombinant human EGF (ImmunoTOOLs) for 5 minutes.

### Transfections

*WNT5A* KD was performed in HDFb using 25μM siRNA targeted to *WNT5A* (si*WNT5A*) (siTOOLs Biotech) for 48 hours with Lipofectamine RNAiMax (Thermo Fisher Scientific). Simultaneously, HDFb were transfected with universal negative control siRNA (siTOOLS Biotech) for 48 hours to generate siNEG HDFb.

### Cellular viability

The cell viability at various concentrations of all pharmacological metabolic drugs was determined using resazurin-based PrestoBlue (Thermo Fisher Scientific). After incubation with PrestoBlue for 30 minutes, cell absorbance at 560nm and 600nm was monitored using a SpectraMax iD3 plate reader (SpectraMax) and the absorbance values were used to analyse cellular viability. HDFb were pre-treated for 2 hours with 3% hydrogen peroxide (H_2_O_2_) (Thermo Fisher Scientific) as a positive control.

### Caspase 3/7 apoptosis quantification

Apoptosis was assessed in HDFb through the quantification of activated caspase-3/7 using the Caspase-Glo® 3/7 Assay System (Promega) according to the manufacturer’s instructions in 96 well plates. Briefly, cultured HDFb were incubated with a 100X stock solution of Caspase-Glo® 3/7 Red Reagent at a 1:100 dilution for 1 hour. images were automatically captured and analysed fluorescence proportional to activated caspase-3/7 relative to cell confluence using the IncuCyte ZOOM system (Sartorius) photographing in phase and red image channels. Red fluorescence units (RFU) were normalized to measured cell confluency for integrated intensity fluorescence units (RFU*µm^2^/image). As a positive control for apoptosis events, a subset of HDFb were treated with 50μM etoposide (Thermo Fisher Scientific).

### Protein extraction and western blotting

Whole cell lysate fractions were isolated from skin biopsies, as well as HDFb as follows: cell pellet obtained was resuspended in RIPA buffer (Thermo Fisher Scientific) and kept at ice for 30 minutes with the inversion at 10-minute intervals. Resuspended cells then were centrifuged at 13000g for 10 minutes and the cell supernatant was subsequently used for western blotting. Isolated subcellular fractions (50 ng) were loaded on 4-20% gradient SDS polyacrylamide gels (Biorad) and ran at 80V followed by 120V for 60-90 minutes. Gel was transferred onto nitrocellulose membrane and blocked in 5% skim milk (Thermo Fisher Scientific) followed by incubation with primary antibody at 4°C overnight with slow shaking. On the next day, membranes were washed with TBST (Tris Buffered Saline with 0.05% Tween20) three times for five minutes, incubated with HRP-conjugated secondary antibody for 1 hour, washed with TBST, and finally detected with femto ECL solution (Thermo Fisher Scientific). The Anti-rabbit or Anti-mouse secondary antibody was used at 1:10000 dilution. The list of antibodies used in this experiment and others is described in **Table 1**.

### Mitophagy flux western blot

Protein levels of BNIP3 (Cell Signaling Technology), NIX (Cell Signaling Technology), and TFAM (Thermo Fisher Scientific) were determined through WB in HDFb in a mitophagy flux assay. In order to inhibit the degradation of lysosomal contents and recycling of mitophagy receptors, HDFb were treated acutely with 50μM CQ for 1 hour prior to lysate harvesting and protein extraction as detailed above. Western blot quantification of proteins was performed as described above.

### Actin polymerisation western blot

The ratio of protein levels of F-actin to G-actin was measured through western blotting as an indicator of actin polymerization. Here, following treatment and transfection, HDFb were rinsed with PBS twice before incubation in lysis buffer containing F-actin stabilising agents (1M HEPES, 1M KCl, 10% Triton-x100, 1M MgCl_2_, and 100mM ATP) for 10 minutes at 37℃, followed by scraping and collection on cell lysate and centrifugation (Optima TLX, Beckman Coulter) at 100,000g for 1 hour at 37℃. Following ultracentrifugation, the supernatant was collected into a tube labelled G-actin, and the pellet was resuspended in 100μl F-actin stabilizing lysis buffer and mechanically digested with magnetic beads. To determine F-actin to G-actin ratio, equal amount of F-actin and G-actin protein (35ng) were subjected to western blotting, as described previously, using an anti-β-actin-peroxidase mouse monoclonal antibody (1:10,000, Sigma Aldrich).

### RT-qPCR

Total RNA was extracted from cells using Trizol reagent (Thermo Fisher Scientific). After determining the sample quality, reverse transcription was performed using the RevertAid First Strand cDNA Synthesis Kit (Thermo Fisher Scientific). Gene expression was quantified by SYBR Green expression assays (Thermo Fisher Scientific) and normalized with β-actin. The primers used in this study is described in **Table 2**.

### RNAseq analysis of publicly-available human wounded tissue

Differential expression (DE) data of mRNA was derived from RNAseq of full-thickness lower-extremity skin biopsies from healthy donors as well as biopsies from non-healing skin of venous ulcer (VU) patients obtained from a publicly-available online tool (http://130.229.28.87/shiny/miRNA_Xulab/) and publication^22^. Adjusted p-values (≤ 0.05) were used to determine genes with significant differential expression and were plotted on a heat map based on log_2_(fold change).

### RNAseq gene expression analysis of UroA-treated HDFb

HDFb treated with either vehicle, 5μM UroA, or 25μM CQ (n = 4 biological repeats per condition) underwent RNA extraction as stated above and RNA quality and quantity was analyzed with Nanodrop 1000, Qubit 4.0 Fluorometer (Life Sciences), and Agilent 5300 Fragment Analyzer (Agilent). RNA from samples were then sent for gene expression profiling using Illumina NovaSeq with Poly(A) selection at the Azenta Life Sciences, Germany and RNA sequencing libraries created using NEB. Next Ultra RNA Library Prep Kit for Illumina (NEB) following manufacturer’s instructions. Briefly, mRNAs were first enriched with Oligo(dT) beads. Enriched mRNAs were fragmented for 15 minutes at 94°C. First strand and second strand cDNAs were subsequently synthesised. cDNA fragments were end repaired and adenylated at 3’ends, and universal adapters were ligated to cDNA fragments, followed by index addition and library enrichment by limited-cycle PCR. Sequencing libraries were validated using NGS Kit on the Agilent 5300 Fragment Analyzer, and quantified by using Qubit 4.0 Fluorometer. The sequencing libraries were multiplexed and loaded on the flowcell on the Illumina NovaSeq X plus instrument according to manufacturer’s instructions. The samples were sequenced using a 2×150 Pair-End (PE) configuration v1.5. Image analysis and base calling were conducted by the NovaSeq Control Software v1.7 on the NovaSeq instrument. Raw sequence data (.bcl files) generated from Illumina NovaSeq was converted into fastq files and de-multiplexed using Illumina bcl2fastq program version 2.20. One mismatch was allowed for index sequence identification. Sequence reads were trimmed to remove possible adapter sequences and nucleotides with poor quality using Trimmomatic v.0.36. The trimmed reads were mapped to the Homo sapiens reference genome available on ENSEMBL using the STAR aligner v.2.5.2b. The STAR aligner is a splice aligner that detects splice junctions and incorporates them to help align the entire read sequences. BAM files were generated as a result of this step. Unique gene hit counts were calculated by using feature Counts from the Subread package v.1.5.2. Only unique reads that fell within exon regions were counted. A PCA analysis was performed using the “plotPCA” function within the DESeq2 R package. The plot shows the samples in a 2D plane spanned by their first two principal components. After extraction of gene hit counts, the gene hit counts table was used for downstream differential expression analysis. Using DESeq2, a comparison of gene expression between the groups of samples was performed. The Wald test was used to generate p-values and Log2 fold changes. Genes showing at least 0.50-fold change with adjusted p-value <0.05 after treatment were used for creating a list of differentially expressed genes for enrichment analysis. Enrichr analysis (https://maayanlab.cloud/Enrichr/) ^62^ was performed to understand the functionally altered pathways in cells treated with UroA compared to vehicle.

### Putative CREB binding site analysis

Putative CREB binding sites within the *WNT5A* promoter region were identified using the JASPAR database^63^. The canonical CREB1 position weight matrix (MA0018.1) was used for scanning the *WNT5A* promoter sequence, which was retrieved for transcript ENST00000264634 from the Eukaryotic Promoter Database (EPD)^64^. Binding sites were identified using a relative profile score threshold of 80%. Predicted motifs were mapped relative to the *WNT5A* transcription start site and annotated by strand orientation.

Visualization of the promoter architecture and binding motifs was performed using R (version 4.5.0, R Core Team, 2024). Genomic plots displaying exon structure, promoter region, transcription start site, and predicted binding sites were generated using the ggplot2^65^ and gggenes packages. Putative binding site sequences were visualized using sequence logos created with the ggseqlogo package^66^ in R.

### Human wounded tissue mitophagy immunofluorescence

Intact, 1 dpw, 7 dpw FFPE skin biopsies (all n = 5) were sectioned at 10μm thickness and stored at 4°C. After incubation for 60 minutes at 65°C, the sections were de-parrafinised and rehydrated as standard. Next, sections were washed in TBST three times for five minutes, followed by antigen retrieval in 10mM sodium citrate pH 6.0 for 7 minutes. Next, sections were blocked with 10% normal goat serum (NGS) (Thermo Fisher Scientific) for 1 hour at room temperature (RT) before another wash cycle. Sections were then incubated with primary antibody for 1 hour at RT (diluted in 10% NGS) and then washed. After washing, sections were incubated with secondary antibody for 1 hour at RT, washed, then incubated with 1μg/ml Hoechst 33342 (Thermo Fisher Scientific) for 10 minutes at RT. Finally, after another wash cycle, sections were mounted on coverslips with EcoMount (Biocare) and stored at 4°C until imaging.

### Live cell immunofluorescence of mitophagy and mitochondrial morphology during *in vitro* wound healing

HDFb were seeded onto μ-dish 35mm with a 4 well insert (Ibidi). When cells were confluent, the inserts were removed using a sterilised tweezer, creating a 500μm gap between cell populations. In order to quantify mitophagy, mitochondrial mass, and mitochondrial morphology at different stages of wound healing, cells were stained with 100nM MitoTracker Green (MTGr), 100nM Tetramethylrhodamine (TMRE), and 60nM LysoTracker Deep Red (LTDR) (all Thermo Fisher Scientific) for 15 minutes. After staining, the dye was washed out with three rinses of Dulbecco’s Phosphate Buffered Saline (PBS) (Thermo Fisher Scientific). Finally, a 100nM TMRE imaging bath was added before the cells were imaged at 63x magnification.

### Live cell cytosolic Ca^2+^ staining and imaging

Cytosolic Ca^2+^ imaging was performed on a Zeiss Airyscan LSM880 confocal microscope in an environmental chamber that maintained the temperature at 37 °C and 5% CO_2_ and imaged at ×20 magnification. HDFb were seeded onto μ-slide eight-well glass bottom plates (Ibidi) and pre-treated with either UroA, rhWNT5A, DFP, or vehicle for 24 h.

HDFb were stained with the ratiometric dye Fura Red AM (5μM in HBSS with 2 mM Ca^2+^ (Thermo Scientific Fisher) + 0.05% Pluronic acid F-127 (Sigma-Aldrich) for 1 hour followed by a 30-min rest at 37 °C. Time-lapse imaging was performed at ×20 magnification using the 488nm laser with 420nm and 480nm excitation wavelengths and 576–638nm and 638–759nm emission filters, with images taken at 10 second intervals. Following the setting of imaging conditions based on unstained cells and Ca^2+^ concentration calibration (Eq. (1)) with 100µM BAPTA-AM (*Rmax*) (R&D Systems) and 1µM ionomycin (*Rmin*), time-lapse imaging of baseline levels was followed by addition of 1µM Tg.

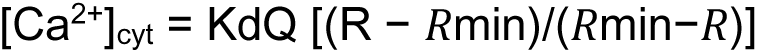

Equation (1) – where R represents the fluorescence intensity ratio Fλ1/Fλ2, in which λ1 and λ2 are the fluorescence detection wavelengths for the ion-bound and ion-free indicator, respectively. Kd is the Ca^2+^ dissociation constant of the indicator. Q is the ratio of Fmin to Fmax at λ2.

### F-actin capping protein immunofluorescence

Immunofluorescence staining of F-actin capping protein was performed on HDFb seeded into μ-slide eight-well glass bottom plates (Ibidi) according to manufacturers instructions. Briefly, following transfection of siNEG and si*WNT5A*, as well as respective treatments, cells were fixed with 4% paraformaldehyde (PFA) (Thermo Fisher Scientific) for 10 minutes before washing with TBS and permeabilisation with 0.1% TritonX-100 (Sigma Aldrich) for 10 minutes. Immediately after, cells were washed with TBS and then blocked with 5% bovine serum albumin (BSA) (Thermo Fisher Scientific) for 1 hour at RT before incubation with primary F-actin capping protein antibody (Thermo Fisher Scientific) for 1 hour at RT, with gentle agitation. Following a wash cycle with TBS, cells were incubated with secondary antibody for 1 hour at RT with gentle agitation. After washing with TBS, cells were then incubated with CellBrite Green (1:100) (Nordic Biosciences) for 10 minutes at RT before incubating with DAPI (1:2000) for 5 minutes at RT, washed with PBS and stored at 4°C.

### Phalloidin immunofluorescence

For fixed cell imaging of phalloidin, HDFb were seeded into μ-slide eight-well glass bottom plates (Ibidi). Following transfection, cells were scratched with a P1000 tip in order to create a manual scratch and migrating cells, and treated for 24 hours with respective treatments. The following day, after 5 minute treatment with 100ng/ml EGF in the EGF-treated HDFb, cells were fixed with 4% PFA (Thermo Fisher Scientific) for 10 minutes, washed with PBS for three cycles of 5 minutes, then permeabilised with 0.1% TritonX-100 (Sigma Aldrich) for 5 minutes. Following another wash cycle with PBS, cells were blocked with 5% BSA (Thermo Fisher Scientific) for 1 hour at RT. Following 1 wash with PBS, cells were incubated with Phalloidin-iFluor 555 reagent (1:1000) (Abcam) for 1 hour at RT. After a wash cycle with PBS, cells were incubated with DAPI (1:2000) for 5 minutes at RT, washed with PBS and stored at 4°C.

### Confocal microscopy and image analysis

IF and live cell imaging was performed on a Zeiss LSM880 with airyscan. For the assessment of mitophagy markers in frozen skin sections, both non-primary control (NPC) and primary ab-stained sections were imaged at x20 magnification. For 1 and 7 dpw, images were taken of the wound site. After subtraction of background fluorescence from the NPC, fluorescence intensity of the respective markers was quantified for each patient and each wound category using Zeiss Zen 3.0 (Blue) software. Following, the fluorescence intensity of each mitophagy marker was normalised to the mitochondrial mass (TOMM20) for each section, then normalised to the TOMM20-normalised fluorescence intensity for the respective intact section.

With regards to the quantification of mitochondrial factors from live cell imaging of HDFb, cells were classed into three categories depending on their proximity to the wound: stationary, which were localised in the main body of cells far away from the wound; wound edge (WE), which were found on the boundary of the wound; and migrating, which were in the gap between the two cell populations. For the quantification of mitophagy, the number of foci of colocalized MTGr, TMRE, and LTDR events was determined for each cell. The Mitochondria Analyzer ImageJ plugin^67^ was used to quantify mitochondrial mass, aspect ratio (AR), and form factor (FF). All images were acquired as Z-stacks at 63x magnification. Three biological repeats with 30 cells for each category were acquired at 0, 24, 48, 72, and 96 hours after removal of the insert.

For imaging of phalloidin-stained HDFb, cells that had migrated into the manual scratch were imaged using the DAPI (nuclei) and Alexa 547 channels (phalloidin) at 63x magnification, with laser strength and gain set following after normalization to NPC. All images were acquired as Z-stacks, with three biological repeats containing 15 cells in each condition. With regards to analysis, phalloidin fluorescence intensity was quantified using the plot profile function after manually drawing a 60μm line from the leading edge towards the nuclei. Measurements of the distance from the leading edge to nuclei was undertaken on Zen Blue (Zeiss).

### Human *ex vivo* full-thickness wound explant model

Full-thickness human skin biopsies were obtained from abdominal reduction surgeries of healthy donors (Nordiska Kliniken, Stockholm) (n = 5) and used as *ex vivo* explant models of human skin wound healing, as previously described^68^. 6mm punch biopsies with 2mm wounds were obtained from four healthy patients, washed in PBS, and cultured in supplemented DMEM with mitophagy inducers and inhibitors at various concentrations. Four explants from each individual patient were used for each respective treatment condition. The samples were maintained at 37°C with 5% CO_2_ for 7 days. At 0, 1, 3, 5, and 7 dpw, the samples were stained with 50μM 5-chloromethylfluorescein diacetate (CMFDA) dye and imaged on a Nikon Eclipse NI-E upright fluorescent microscope (Nikon). The area of the wound at each time point was quantified and normalised against the initial wound area in order to quantify wound healing rate percentage.

### Aged mouse full-thickness wound model

In order to study the effect of UroA treatment on wound healing *in vivo*, we employed aged female C57BL/6 mice (> 70 weeks), supplied from Janvier Labs, France, and housed at the animal facilities of the Zone d’Evaluation Fonctionnelle (ZEF), France. Protocols were assessed and approved by Ethical Committee N°CEEA-122, regularly assessed for their well-being, and fed *ad libitum* diet.

After a 5 day acclimitisation period, two 6mm full-thickness biopsies were taken from each animal (n = 24) following shaving of the back under isoflurane anesthesia, and wounds were immediately splinted with silicone rings and allowed to heal under moist conditions. Splints were changed every 2 days, and anti-inflammatory (carprofen) was administered for 2 days daily. Of the 24 animals, 12 were administered daily with 25mg/ml UroA (BOC Sciences) orally through gavage, whilst 12 were administered vehicle (0.5% w/v carboxymethylcellulose, 0.25% v/v; Tween 80 in dH_2_O), with wounds imaged on days 0, 2, 4, 6, 8, 10, 12, 14, and 15 post wounding. At 2 timepoints (5 and 15 dpw), wounds were harvested and either stored in 4% PFA, or immediately snap-frozen and placed in dry ice for histology staining and imaging. Following harvesting of wounds, animals were euthanised. Wound closure (%) was calculated from images as wound area_time point_/wound area_0 dpw_ x 100%, whilst re-epithelialisation was calculated as neo-epidermis_time point_/neo-epidermis_0 dpw_ x 100%.

### Histochemistry staining and imaging

Following sectioning of FFPE blocks, 5 and 15 dpw biopsies from aged mouse wound models were stained by haematoxylin and eosin (H&E) histochemistry (Klinisk Patologi, Akademiska Sjukhuset, Uppsala, Sweden) as standard and imaged on ZEISS Axioscan 7 scanner (FENO, Karolinska Institutet, Sweden) at 20x magnification. Neo-epidermis length (μm) and epidermal thickness (μm) was calculated for each sample using ZEISS Zen Blue software. Additionally, the number of newly formed hair follicles residing in the wound matrix was quantified and then normalised to wound area (μm).

Masson’s trichrome staining was undertaken in order to quantify collagen deposition within wound beds. Here, 5 and 15 dpw sections were subjected to Masson’s trichome histochemistry (Klinisk Patologi, Akademiska Sjukhuset, Uppsala, Sweden) as standard and imaged using a Nikon Eclipse NI-E upright fluorescent microscope at 4x and 20x magnification. Collagen deposition within the wound bed was quantified following colour deconvolution using the Colour Deconvolution 2 plugin on ImageJ, followed by thresholding, in which the percentage of area covered with collagen (blue) was quantified for each section.

### Scratch wound healing assay

HDFb were seeded on 96-well ImageLock plates (Sartorius). After the cells reached monolayer confluence they were scratched with the Wound Maker 96 (Essen Bioscience) in order to generate wounds. In addition, where stated, cells were treated with and without a 2 hour pre-treatment with 10μg/ml MMC (Sigma Aldrich) prior to scratching, in order to inhibit cell proliferation. After scratching, the cells were incubated in an IncucyteZOOM (Sartorius) and imaged every 2 hours at 10x magnification for 48-72 hours. Relative wound density (%) was quantified on Incucyte 2022A software (Sartorius).

### XF bioenergetic assay

HDFb cells were seeded on XFe24-well cell culture microplates (Agilent) at 2.8−3.5 × 10^4^ cells/well for 16−24 hours. OCR (percentage oxygen consumption rate over baseline) and ECAR (percentage extracellular acidification rate over baseline) were measured using an XFe24 extracellular flux analyser (Agilent) in a minimal DMEM assay medium supplemented with 10 mM glucose (Thermo Fisher Scientific), 1 mM sodium pyruvate (Thermo Fisher Scientific), and 2 mM L-glutamine (Thermo Fisher Scientific) for 45−60 minutes for equilibration. To determine the acute responses to pharmacological inhibitors and enhancers of UroA and CQ, measurements were taken before and after exposure to compound injections at regular intervals for 3 cycles. Data was analysed using the Wave Software (Agilent) and expressed as a percentage of oxygen consumption rate or extracellular acidification rate change over the pre-exposure baseline rate.

### Statistical Analysis

Statistical significance was determined by two tailed Student’s t-test. The significance among two group was determined by paired or unpaired t-tests, and by Two-way ANOVA for multiple groups, both on GraphPad Prism Version 6. The p-values of GSEA analysis were calculated by using Fisher’s exact test. P-value < 0.05 was determined to be statistically significant.

## Data availability

Data sequenced in this study have been deposited in GEO under accession numbers: PRJNA1327083.

Any additional information required to reanalyse the data reported in this paper is available from the lead contact upon request.

## Acknowledgments

We thank the patients for participating in this study and providing the biopsies as well as research nurse Helena Griehsel. This work was supported by grants from Hudfonden, Swedish Science Council, Swedish Society for Medical Research, Åke Wibergs Stiftelse, LEO foundation, ALF Medicin Stockholm, Jeanssons Stiftelse, Wallenberg Foundation, Tore Nilssons Stiftelse, and Stiftelsen Sigurd and Elsa Goljes Minne.

## Author contributions

MH, JW, and EBW conceptualised the project and planned the experiments. MH, MT, NW, MC, SH, GAQ, and EBW performed experiments and analysed data. All authors contributed to the manuscript writing and editing.

## Resource availability

### Lead contact

Further information and requests for resources and reagents should be directed to and will be fulfilled by the lead contact, Jakob Wikström.

## Materials availability

This study did not generate new, unique reagents.

---

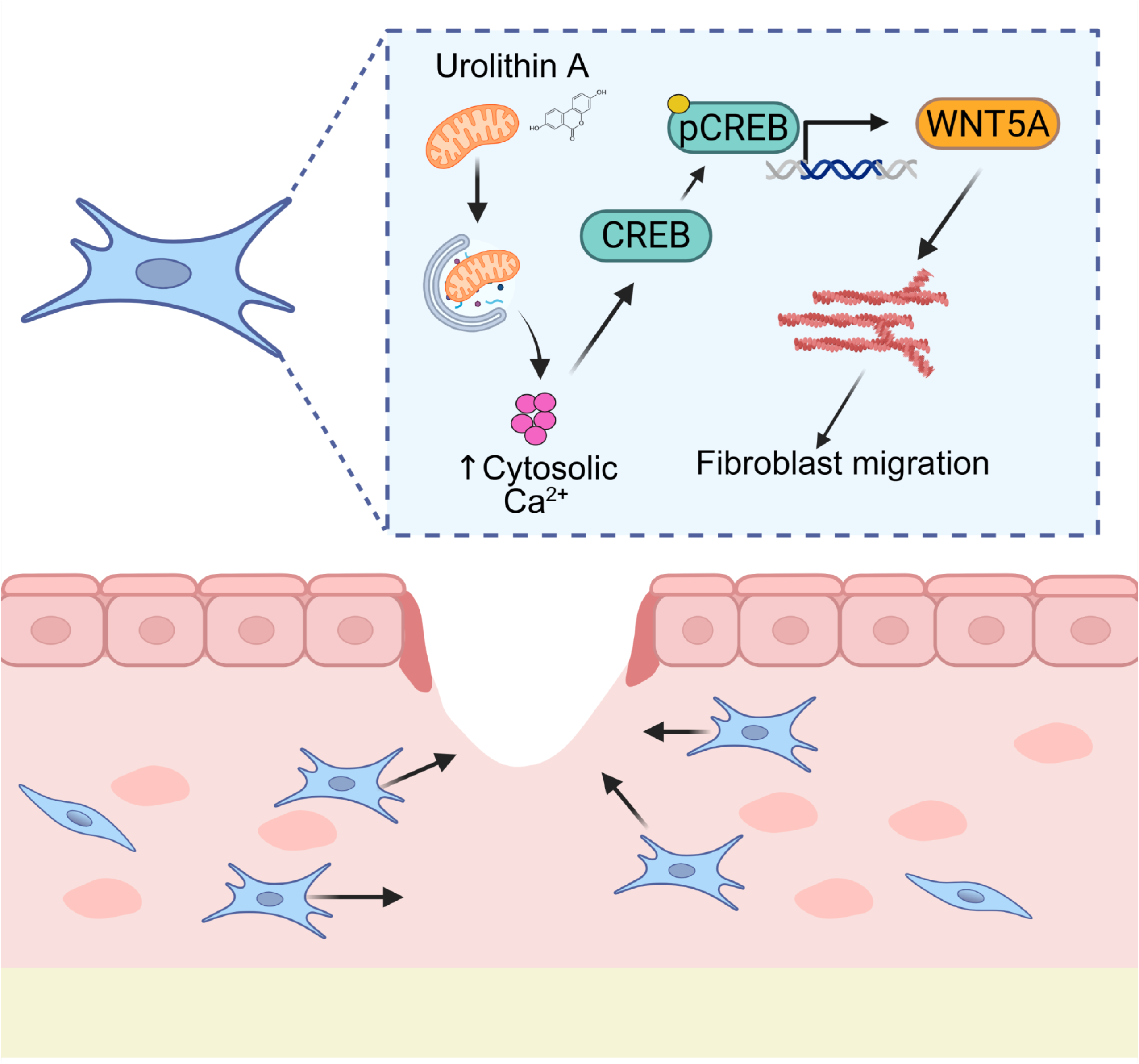

**Figure S1.**
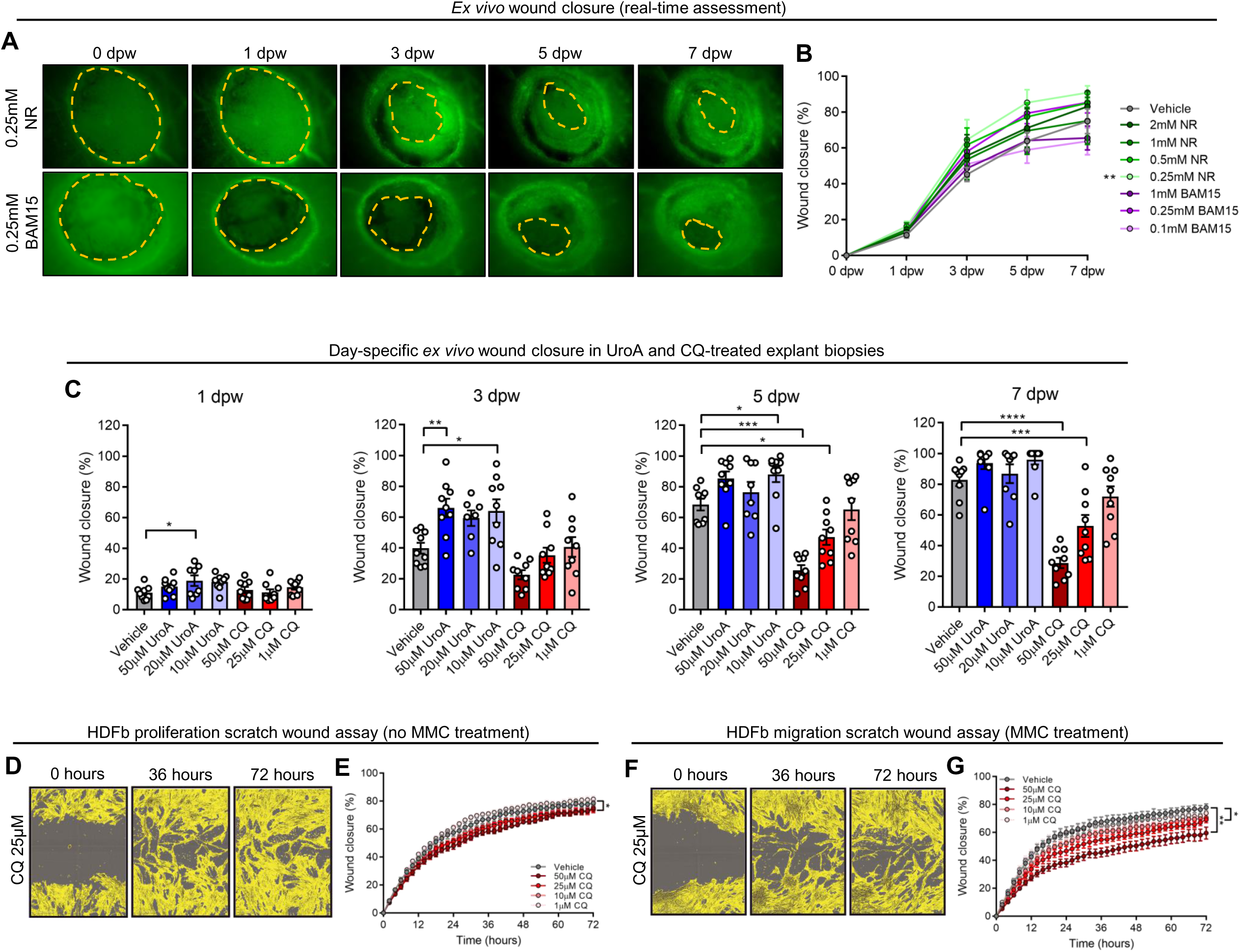
Mitophagy accelerates and CQ inhibits re-epethelialisation and HDFb migration, related to. **Figure 3**. (**A**) Representative fluorescence images of wounded explant *ex vivo* biopsies treated with vehicle, 0.25mM NR and 0.25mM BAM15 at 0, 1, 3, 5, and 7 dpw. Scale bar = 200μm. (**B**) Quantification (mean ± SEM) of wound healing time of wounds treated with various concentrations of UroA and CQ. Two-way ANOVA,; *** = p < 0.001. N = 5 biological replicates. (**C**) Graphs (mean ± SEM) of wound closure % of vehicle, UroA, and CQ treated *ex vivo* full-thickness explants at 1, 3, 5, and 7 dpw. Two-way ANOVA,; **** = p < 0.0001, *** = p < 0.001, ** = p < 0.01, * = p < 0.05. (**D-E**) Representative images and (**E**) quantification (mean ± SEM) of scratch wound assay in HDFb treated with vehicle and CQ. (**F-G**) Representative images and (**G**) quantification (mean ± SEM) of scratch wound assay in HDFb treated with vehicle + MMC and CQ + MMC over 72 hours. Two-way ANOVA, ** = p < 0.01, * = p < 0.05. N = 3 biological replicates containing at least 3 technical replicates.

**Figure S2.**
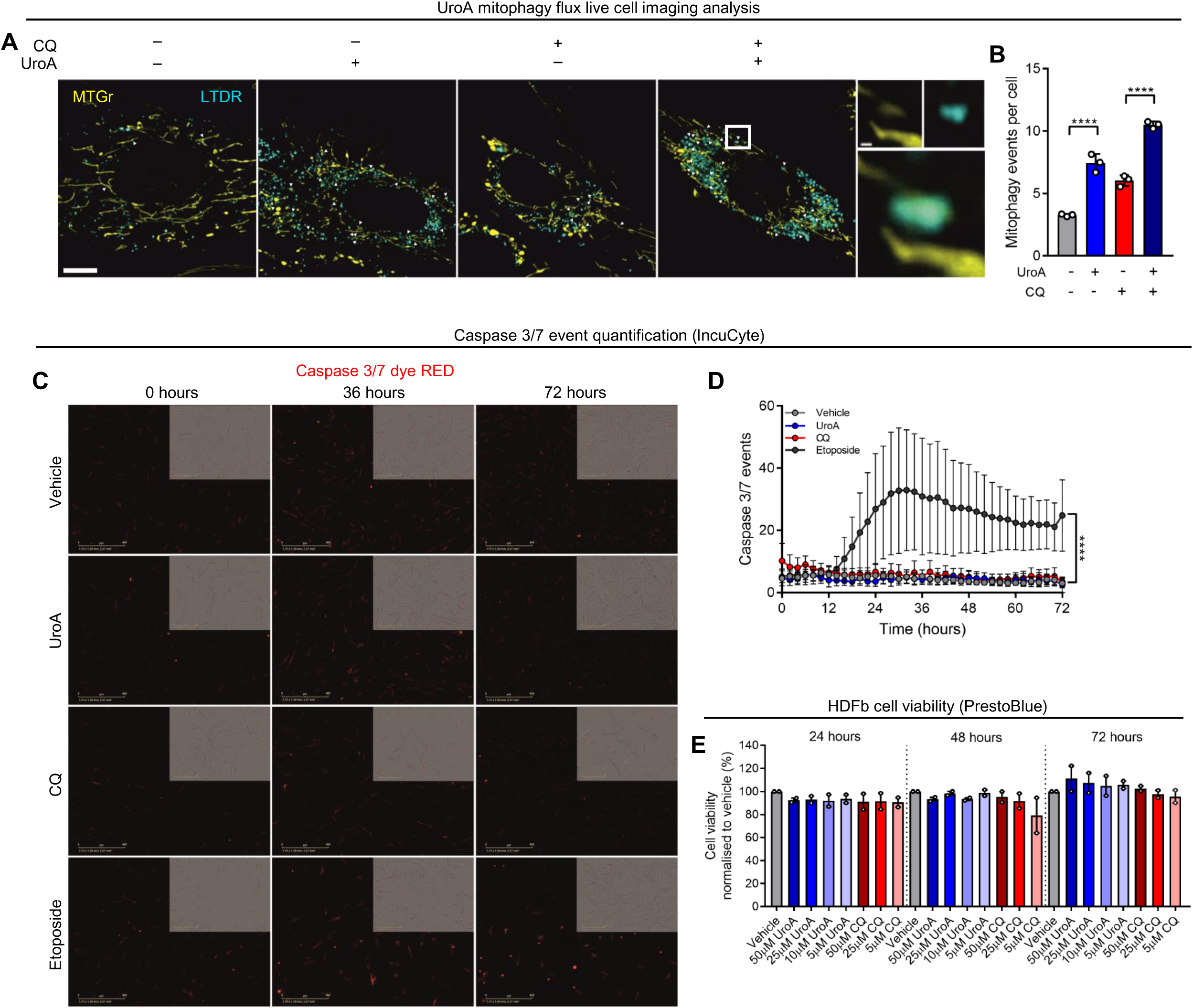
UroA increases mitophagy and is not toxic in HDFb, related to. **Figure 3**. (**A-B**) Representative fluorescence images and (**B**) quantification (mean ± SEM) of mitophagy events per cell in HDFb following staining with MitoTracker Green and LysoTracker Deep Red. Two-way ANOVA,; **** = p < 0.0001. N = 3 biological repeats containing at least 15 cells. (**C-D**) Representative fluorescence images and (**D**) quantification (mean ± SEM) of Caspase 3/7 events over 72 hours in HDFb stained with Caspase 3/7 dye RED. Two-way ANOVA,; **** = p < 0.0001. N = 3 biological repeats. (**E**) Quantification (mean ± SEM) of percentage cell viability normalised to vehicle in HDFb treated with various concentrations of UroA and CQ over 72 hours.

**Figure S3.**
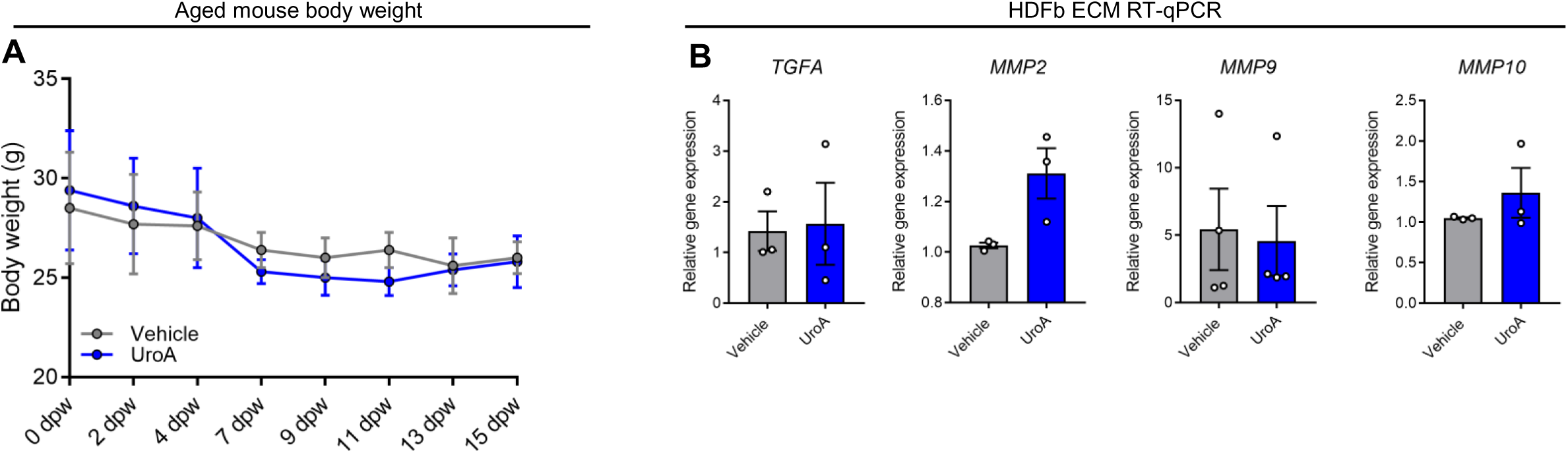
No change in mouse body weight and ECM gene expression in HDFb, related to. **Figure 4**. (**A**) Quantification (mean ± SEM) of body weight (g) of mice treated with either vehicle or UroA for 15 days. Two-way ANOVA. (N = 25 mice). (**B**) Quantification (mean ± SEM) of gene expression of ECM-related related genes in HDFb treated with either vehicle for UroA for 24 hours. Student’s t-test. Each dot represents an individual biological repeat containing at least 2 technical repeats.

**Figure S4.**
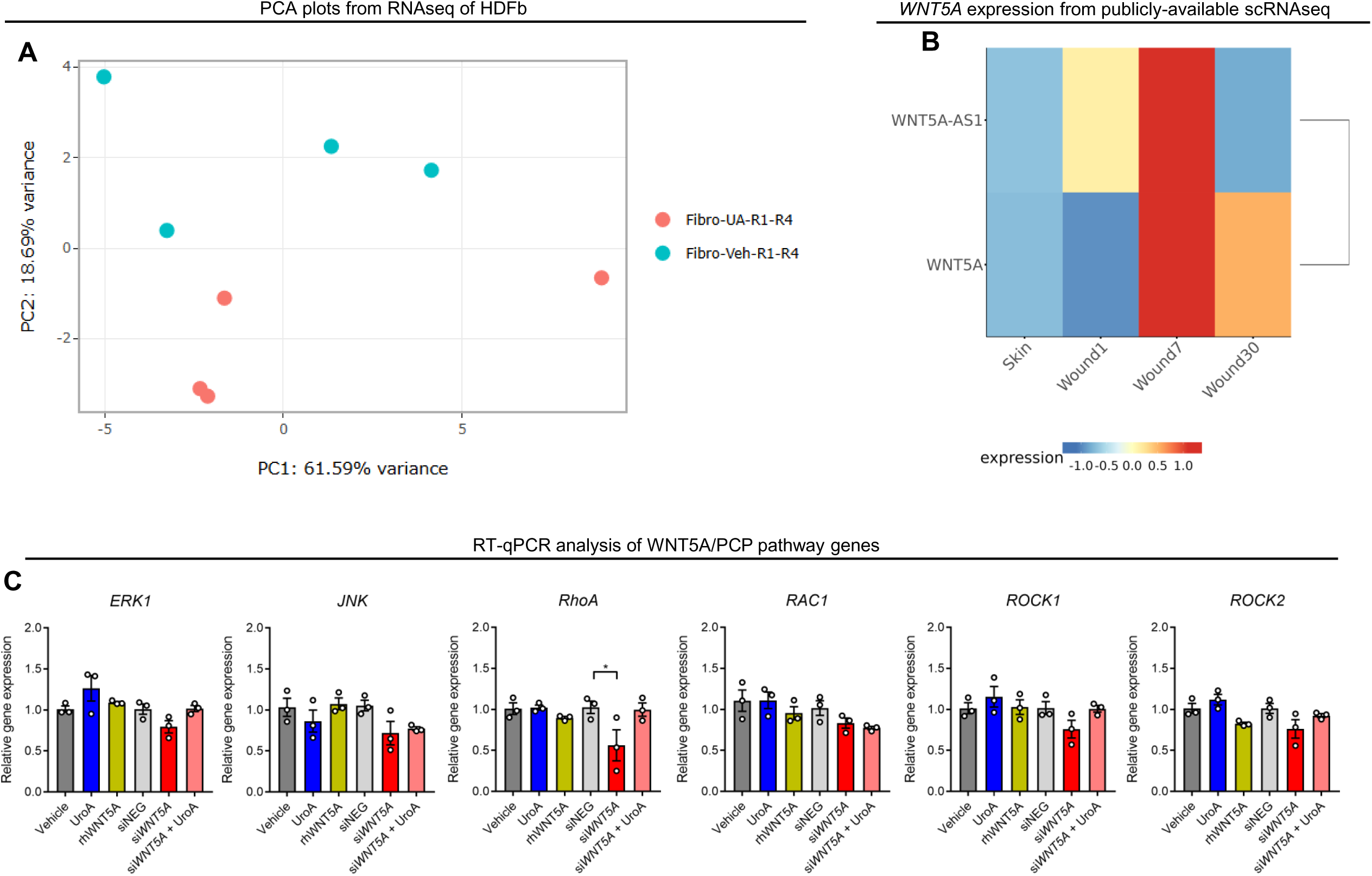
***WNT5A* is increased in fibroblasts in human wound healing samples and UroA does not increase WNT5A/PCP pathway genes in HDFb, related to Figure 5**. (**A**) PCA showing similarity analysis of Vehicle and UroA-treated HDFb used for RNAseq. (**B**) Heatmap showing *WNT5A* and *WNT5A-AS1* expression in FB-1, FB-II, FB-III, and FB-prolif clusters from scRNAseq data, derived from https://shiny.xulandenlab.com/shiny/scwoundatlas/. (**C**) Quantification (mean ± SEM) of the relative gene expression of genes associated with the WNT5A/PCP pathway in HDFb. Two-way ANOVA, * = p < 0.05. N = 3 biological replicates containing at least 2 technical repeats.

**Figure S5.**
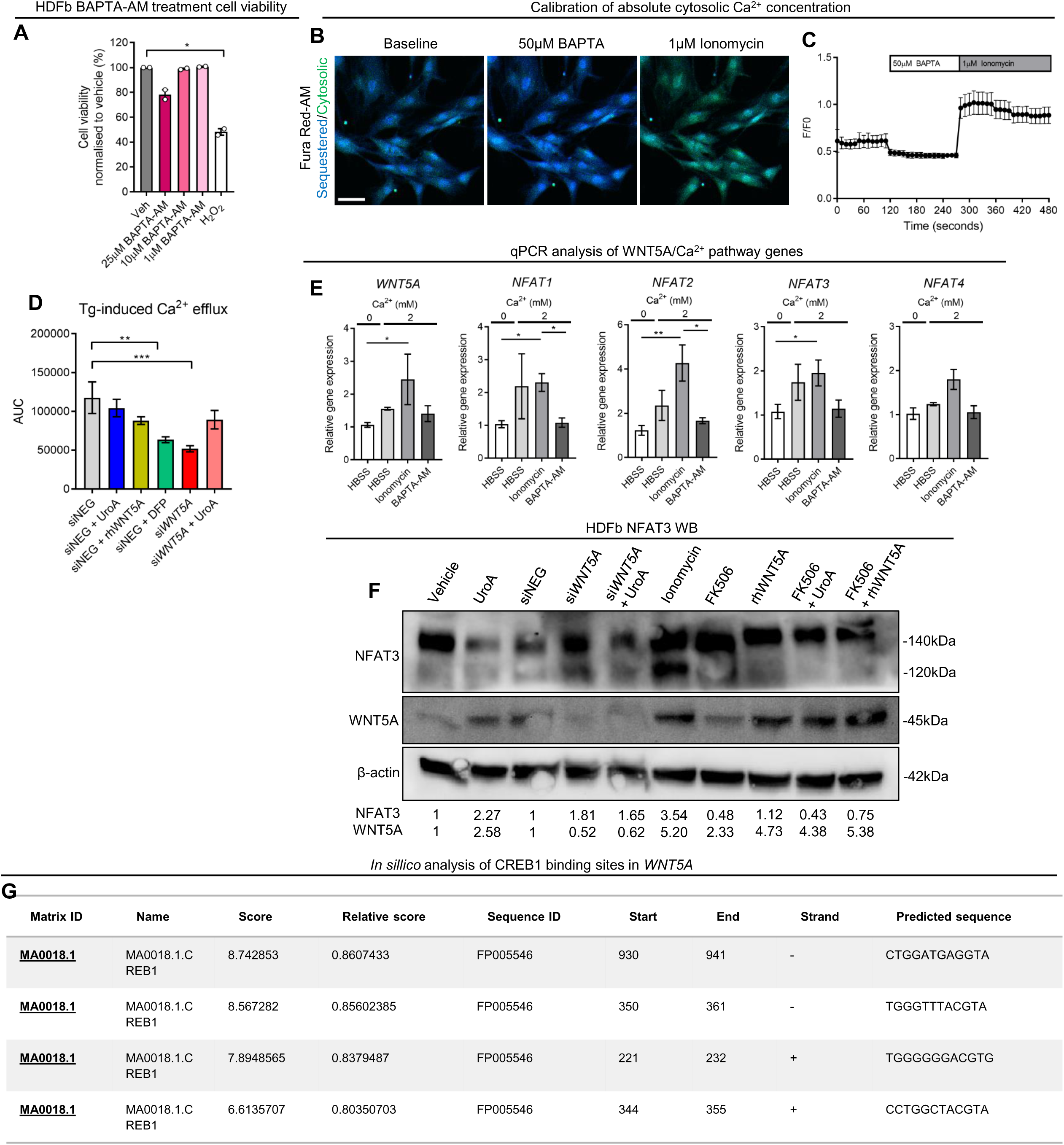
**Ca2+ signalling increases WNT5A transcription in HDFb, related to Figure 6.** (**A**) Quantification (mean ± SEM) of cell viability in HDFb treated with various concentrations of Ca2+ quencher BAPTA-AM. Two-way ANOVA, * = p < 0.05. Each dot represents an individual biological repeat (N = 2) containing at least 3 technical repeats. (**B-C**) Representative fluorescence images and (C) graph of F/F0 ratio of HDFb stained with Fura Red-AM at baseline, after 50µM BAPTA-AM (Rmin), and 1µM ionomycin (Rmax). (**D**) Quantification (mean ± SEM) of area under the curve after Tg-induced Ca2+ efflux in HDFb stained with Fura-Red AM. Two-way ANOVA, *** = p < 0.001, ** = p < 0.01. N = 3 biological repeats containing at least 20 cells. (**E**) Quantification (mean ± SEM) of WNT5A and NFAT gene expression in response to Ca2+ modulation in HDFb. N = 3 biological repeats containing at least 2 technical repeats. Two-way ANOVA, * = p < 0.05. (**F**) Representative immunoblot and relative band expression of NFAT and WNT5A protein levels in HDFb. (G) Motif analysis of putative CREB binding sites in the WNT5A promoter region.

**Figure S6.**
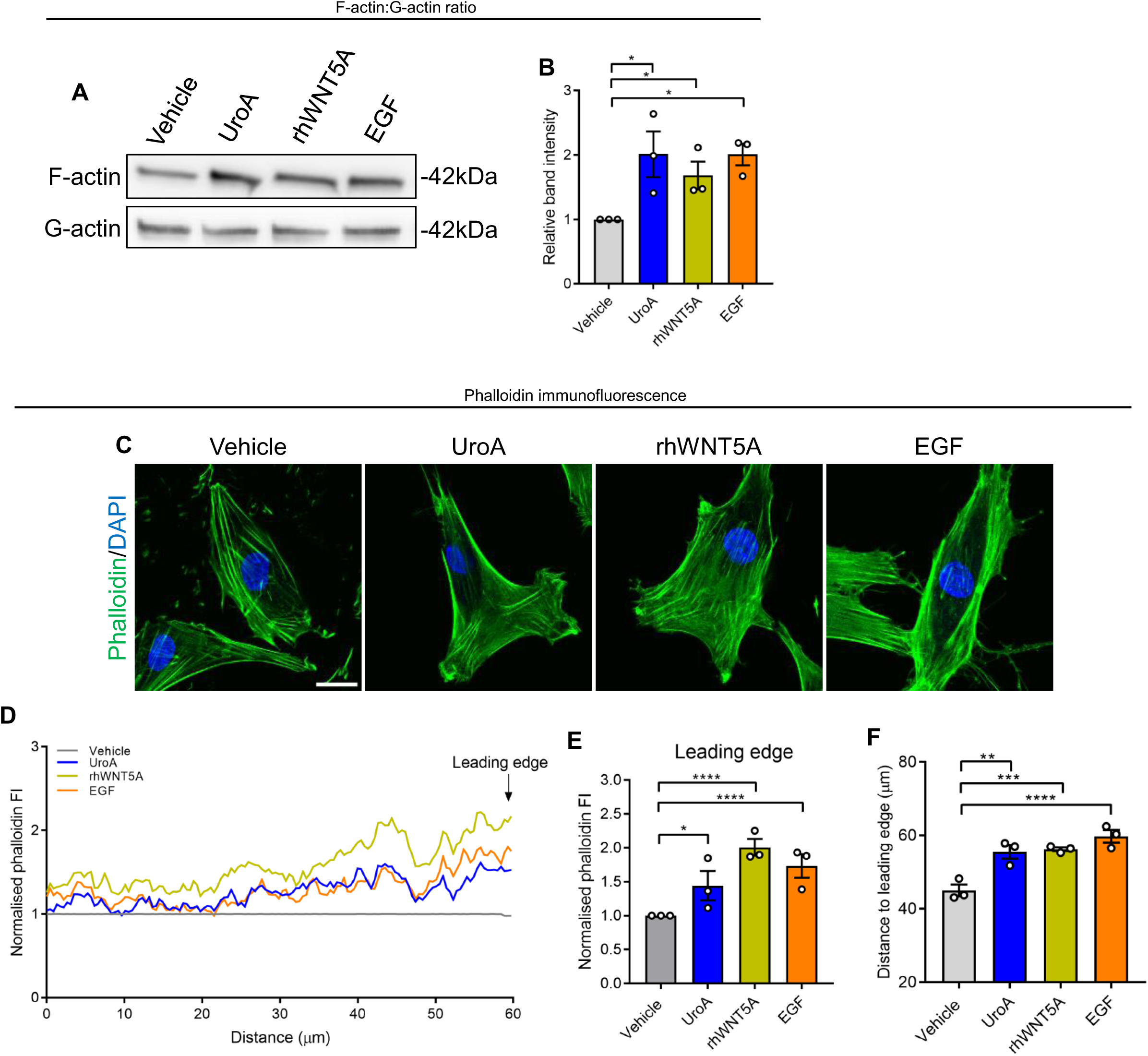
UroA, rhWNT5A, and EGF increase actin polymerisation in HDFb, related to. **Figure 7**. (**A-B**) Representative immunoblot and (**B**) quantification of F-actin and G-actin in HDFb. Each dot represents an biological repeat (n = 3). Two-way ANOVA, * = p < 0.05. (**C**) Representative immunofluorescence images of phalloidin staining in HDFb. Scale bar = 10µm. (**D**) Point plot of phalloidin fluorescence intensity normalised to Vehicle in HDFb. Each line represents the mean of 3 biological replicated each containing 15 cells. 60μm indicates the cell periphery. (**E**) Quantification (mean ± SEM) of the average phalloidin fluorescence intensity at the final 10μm of the cell periphery. Two-way ANOVA, **** = p < 0.0001, * = p < 0.05. Each dot represents an individual biological replicate containing 15 individual cells. (F) Quantification (mean ± SEM) of the distance (μm) from the nuclei to the periphery of the leading edge within HDFb. Two-way ANOVA, **** = p < 0.0001, *** = p < 0.001, ** = p < 0.01. Each dot represents an individual biological replicate containing 15 individual cells.

## References

1. Olsson, M., Järbrink, K., Divakar, U., Bajpai, R., Upton, Z., Schmidtchen, A., and Car, J. (2019). The humanistic and economic burden of chronic wounds: A systematic review. Wound Rep Reg 27 114–125.

2. Järbrink, K., Ni, G., and Sönnergren, H., et al. (2017). The humanistic and economic burden of chronic wounds: a protocol for a systematic review. Syst Rev 6 (15 ).

3. Falanga, V., Isseroff, R., and Soulika, A., et al. (2022). Chronic wounds. Nat Rev Dis Primers 8, 50

4. Torres, M., Silberberg, G., Vegvari, A., Zubarev, R., Hunt, M., Bansal, R., Bachar-Wikstrom, E., and Wikstrom, J. (2024). The temporal dynamics of proteins in aged skin wound healing and comparison to gene expression. Jouornal Of Investigative Dermatology In press.

5. Knoedler, S., Broichhausen, S., Guo, R., Dai, R., Knoedler, L., Kauke-Navarro, M., Diatta, F., Pomahac, B., Machens, H., Jiang, D., and Rinkevich, Y. (2023). Fibroblasts - the cellular choreographers of wound healing. Front Immunol 14:1233800.

6. Eming, S., Martin, P., and Tomic-Canic, M. (2014). Wound repair and regeneration: mechanisms, signaling, and translation. Sci. Transl. Med 6 265sr266– 265sr266.

7. Manchanda, M., Torres, M., Inuossa, F., Bansal, R., Kumar, R., Hunt, M., Wheelock, C., Bachar-Wikström, E., and Wikström, J. (2023). Metabolic Reprogramming and Reliance in Human Skin Wound Healing. J Invest Dermatol S0022-202X(23) 01975-01979.

8. Willenborg, S., Sanin, D., Jais, A., Ding, X., Ulas, T., and Nuchel, J., et al. (2020). Mitochondrial metabolism coordinates stage-specific repair processes in macrophages during wound healing. Cell Metab.

9. Schiffmann, L., Werthenbach, J., and Heintges-Kleinhofer, F., et al. (2020). Mitochondrial respiration controls neoangiogenesis during wound healing and tumour growth. Nat Commun 11(1) (3653 ).

10. Ma, K., et al. (2020). Mitophagy, Mitochondrial Homeostasis, and Cell Fate. Frontiers in Cell and Developmental Biology 8 (467).

11. Levine, B., and Kroemer, G. (2008). Autophagy in the pathogenesis of disease. Cell 132(1), 27–42

12. Doblado, L., Lueck, C., Rey, C., Samhan-Arias, A., Prieto, I., Stacchiotti, A., and Monsalve, M. (2021). Mitophagy in Human Diseases. Int J Mol Sci 22(8) (3903 ).

13. Ganley, I., and Simonsen, A. (2022). Diversity of mitophagy pathways at a glance. J Cell Sci 1 December 2022; 135 (23): jcs259748 135 (23) (jcs259748 ).

14. Hunt, M., Torres, M., Bachar-Wikström, E., and Wikström, J. (2023). Multifaceted roles of mitochondria in wound healing and chronic wound pathogenesis. Frontiers in Cell and Developmental Biology 11.

15. Feng, Y., Li, L., Zhang, Q., He, Y., Huang, Y., Zhang, J., Zhang, D., Huang, Y., Lei, X., Hu, J., and Luo, G. (2023). Mitophagy associated self-degradation of phosphorylated MAP4 guarantees the migration and proliferation responses of keratinocytes to hypoxia Cell Death Discov. 9(1) 168.

16. Feng, Z., Chen, J., Yuan, P., Ji, Z., Tao, S., Zheng, L., Wei, X., Zheng, Z., Zheng, B., Chen, B., et al. (2022). Urolithin A Promotes Angiogenesis and Tissue Regeneration in a Full-Thickness Cutaneous Wound Model. Front Pharmacol. 13:806284.

17. Zhang, J., Zhang, C., and Jiang, X., et al. (2019). Involvement of autophagy in hypoxia-BNIP3 signaling to promote epidermal keratinocyte migration. Cell Death Dis 10, 234

18. Zhang, J., et al. (2017). BNIP3 promotes the motility and migration of keratinocyte under hypoxia. Exp. Dermatol 26, 416–422.

19. Moriyama, M., Moriyama, H., and Uda, J., et al. (2017). BNIP3 upregulation via stimulation of ERK and JNK activity is required for the protection of keratinocytes from UVB-induced apoptosis. Cell Death Dis 8, e2576

20. Moriyama, M., Moriyama, H., Uda, J., Matsuyama, A., Osawa, M., and Hayakawa, T. (2014). BNIP3 plays crucial roles in the differentiation and maintenance of epidermal keratinocytes. J Invest Dermatol 134(6), 1627–1635

21. Simpson, C., Tokito, M., Uppala, R., Sarkar, M., Gudjonsson, J., and Holzbaur, E. (2021). NIX initiates mitochondrial fragmentation via DRP1 to drive epidermal differentiation. Cell Reports 34, 108689.

22. Liu, Z., Zhang, L., and Toma, M., et al. (2022). Integrative small and long RNA omics analysis of human healing and nonhealing wounds discovers cooperating microRNAs as therapeutic targets. eLife 11, e80322

23. Liu, Z., Bian, X., and Luo, L., et al. (2025). Spatiotemporal single-cell roadmap of human skin wound healing. Cell Stem Cell 32(3):479-498.e8.

24. Zimmermann, A., Madeo, F., Diwan, A., Sadoshima, J., Sedej, S., Kroemer, G., and Abdellatif, M. (2024). Metabolic control of mitophagy. Eur J Clin Invest 54(4):e14138.

25. Krantz, S., Kim, Y., Srivastava, S., Leasure, J., Toth, P., Marsboom, G., and Rehman, J. (2021). Mitophagy mediates metabolic reprogramming of induced pluripotent stem cells undergoing endothelial differentiation. J Biol Chem 297(6):101410.

26. Wang, S., Long, H., and Hou, L., et al. (2023). The mitophagy pathway and its implications in human diseases. Sig Transduct Target Ther 8 , 304.

27. Yang, M., Wei, X., Yi, X., and Jiang, D. (2024). Mitophagy-related regulated cell death: molecular mechanisms and disease implications. Cell Death Dis 15(7):505.

28. Pidgeon, R., Mitchell, S., Shamash, M., Suleiman, L., Dridi, L., Maurice, C., and Castagner, B. (2025). Diet-derived urolithin A is produced by a dehydroxylase encoded by human gut Enterocloster species Nat Commun 16(1):999.

29. Andreux, P., Blanco-Bose, W., Ryu, D., Burdet, F., Ibberson, M., Aebischer, P., Auwerx, J., Singh, A., and Rinsch, C. (2019). The mitophagy activator urolithin A is safe and induces a molecular signature of improved mitochondrial and cellular health in humans. Nat Metab 1(6):595-603.

30. Kuerec, A., Lim, X., Khoo, A., Sandalova, E., Guan, L., Feng, L., and Maier, A. (2024). Targeting aging with urolithin A in humans: A systematic review. Ageing Res Rev 100:102406.

31. D’Amico , D., Fouassier , A., Faitg , J., Hennighausen , N., Brandt , M., Konstantopoulos , D., Rinsch , C., and Singh, A. (2023). Topical application of Urolithin A slows intrinsic skin aging and protects from UVB-mediated photodamage: Findings from Randomized Clinical Trials. medRxiv 2023.06.16.23291378

32. Liu, S., Faitg, J., and Tissot, C., et al. (2025). Urolithin A provides cardioprotection and mitochondrial quality enhancement preclinically and improves human cardiovascular health biomarkers. iScience 28 ; 111814.

33. Ryu, D., Mouchiroud, L., and Andreux, P., et al. (2016). Urolithin A induces mitophagy and prolongs lifespan in C. elegans and increases muscle function in rodents. Nat Med 22(8):879-88.

34. Luan, P., D’Amico, D., and Andreux, P., et al. (2021). Urolithin A improves muscle function by inducing mitophagy in muscular dystrophy. Sci Transl Med 13(588):eabb0319.

35. Bansal, R., Torres, M., Hunt, M., Wang, N., Chatzopoulou, M., Manchanda, M., Taddeo, E., Shu, C., Shirihai, O., Bachar-Wikstrom, E., and Wikstrom, J. (2024). Role of the mitochondrial protein cyclophilin D in skin wound healing and collagen secretion. JCI Insight 9(9) e169213.

36. Qin, K., Yu, M., and Fan, J., et al. (2023). Canonical and noncanonical Wnt signaling: Multilayered mediators, signaling mechanisms and major signaling crosstalk. Genes Dis 11(1):103-134.

37. Lee, H., Kim, S., Kwon, N., Jo, S., Kwon, O., and Kim, J. (2024). Single-Cell and Spatial Transcriptome Analysis of Dermal Fibroblast Development in Perinatal Mouse Skin: Dynamic Lineage Differentiation and Key Driver Genes. J Invest Dermatol 144(6):1238-1250.e11.

38. Reddy, S., Andl, T., Bagasra, A., Lu, M., Epstein, D., Morrisey, E., and Millar, S. (2001). Characterization of Wnt gene expression in developing and postnatal hair follicles and identification of Wnt5a as a target of Sonic hedgehog in hair follicle morphogenesis. Mech Dev 107:69–82.

39. Fathke, C., Wilson, L., Shah, K., Kim, B., Hocking, A., Moon, R., and Isik, F. (2006). Wnt signaling induces epithelial differentiation during cutaneous wound healing. BMC Cell Biol 7:4

40. Trinh-Minh, T., Chen, C., Tran, M.C., and al., e. (2024). Noncanonical WNT5A controls the activation of latent TGF-β to drive fibroblast activation and tissue fibrosis. J Clin Invest 134(10):e159884.

41. Shah, R., Amador, C., and Poe, A., et al. (2025). Identification of Wnt-5a Receptors Important in Diabetic and Non-Diabetic Corneal Epithelial Wound Healing. Invest Ophthalmol Vis Sci 66(2):64.

42. Clark, C., Nourse, C., and Cooper, H. (2012). The tangled web of non-canonical Wnt signalling in neural migration. Neurosignals 20(3):202-20.

43. Borbolis, F., Ploumi, C., and Palikaras, K. (2025). Calcium-mediated regulation of mitophagy: implications in neurodegenerative diseases. NPJ Metab Health Dis 3(1):4

44. Berridge, M., Bootman, M., and Roderick, H. (2003). Calcium signalling: dynamics, homeostasis and remodelling. Nat Rev Mol Cell Biol 4(7):517-29.

45. Jiang, D., Christ, S., and Correa-Gallegos, D., et al. (2020). Injury triggers fascia fibroblast collective cell migration to drive scar formation through N-cadherin Nat Commun 11(1):5653.

46. Garbincius, J., and Elrod, J. (2022). Mitochondrial calcium exchange in physiology and disease. Physiol Rev 102(2):893-992.

47. Grossmann, D., Berenguer-Escuder, C., and Bellet, M., et al. (2019). Mutations in RHOT1 Disrupt Endoplasmic Reticulum-Mitochondria Contact Sites Interfering with Calcium Homeostasis and Mitochondrial Dynamics in Parkinson’s Disease. Antioxid Redox Signal 31(16):1213-1234.

48. Szabadkai, G., Bianchi, K., Várnai, P., De Stefani, D., Wieckowski, M., Cavagna, D., Nagy, A., Balla, T., and Rizzuto, R. (2006). Chaperone-mediated coupling of endoplasmic reticulum and mitochondrial Ca2+ channels. J Cell Biol 175(6):901-11.

49. Sassano, M., Felipe-Abrio, B., and Agostinis, P. (2022). ER-mitochondria contact sites; a multifaceted factory for Ca 2+ signaling and lipid transport. Front Cell Dev Biol 10:988014.

50. Wang, H., Xu, J., Lazarovici, P., Quirion, R., and Zheng, W. (2018). cAMP Response Element-Binding Protein (CREB): A Possible Signaling Molecule Link in the Pathophysiology of Schizophrenia. Front Mol Neurosci 11:255.

51. Dyson, H., and Wright, P. (2016). Role of Intrinsic Protein Disorder in the Function and Interactions of the Transcriptional Coactivators CREB-binding Protein (CBP) and p300. J Biol Chem 291(13):6714-22.

52. Wei, H., Deng, M., Ding, R., Wei, L., and Yuan, H. (2024). Macrophage β2-AR activation amplifies inflammation in wound healing by upregulating Trem1 via the cAMP/PKA/CREB pathway Int Immunopharmacol 128:111463.

53. Sakamoto, K., Karelina, K., and Obrietan, K. (2011). CREB: a multifaceted regulator of neuronal plasticity and protection. J Neurochem 116(1):1-9.

54. Zhao, W., Hylton, N., Wang, J., Chindavong, P., Alural, B., Kurtser, I., Subramanian, A., Mazitschek, R., Perlis, R., and Haggarty, S. (2019). Activation of WNT and CREB signaling pathways in human neuronal cells in response to the Omega-3 fatty acid docosahexaenoic acid (DHA). Mol Cell Neurosci 99:103386.

55. Trinh, A., Kim, S., Chang, H., Mastrocola, A., and Tibbetts, R. (2013). Cyclin-dependent kinase 1-dependent phosphorylation of cAMP response element-binding protein decreases chromatin occupancy. J Biol Chem 288(33):23765-75.

56. Wen, A., Sakamoto, K., and Miller, L. (2010). The role of the transcription factor CREB in immune function. J Immunol 185(11):6413-9.

57. Pollard, T. (2016). Actin and Actin-Binding Proteins. Cold Spring Harb Perspect Biol 8(8):a018226.

58. Fung, T., Chakrabarti, R., and Higgs, H. (2023). The multiple links between actin and mitochondria Nat Rev Mol Cell Biol 24(9):651-667.

59. Krause, M., and Gautreau, A. (2014). Steering cell migration: lamellipodium dynamics and the regulation of directional persistence. Nat Rev Mol Cell Biol 15(9):577-90.

60. Edwards, M., Zwolak, A., Schafer, D., Sept, D., Dominguez, R., and Cooper, J. (2014). Capping protein regulators fine-tune actin assembly dynamics. Nat Rev Mol Cell Biol 15(10):677-89.

61. Bachar-Wikstrom, E., Manchanda, M., Bansal, R., Karlsson, M., Kelly-Pettersson, P., Skoldenberg, O., and Wikstrom, J.D. (2021). Endoplasmic reticulum stress in human chronic wound healing: Rescue by 4-phenylbutyrate. Int Wound J 18, 49–61. 10.1111/iwj.13525.

62. Kuleshov, M.V., Jones, M.R., Rouillard, A.D., Fernandez, N.F., Duan, Q., Wang, Z., Koplev, S., Jenkins, S.L., Jagodnik, K.M., Lachmann, A., et al. (2016). Enrichr: a comprehensive gene set enrichment analysis web server 2016 update. Nucleic Acids Res 44, W90–97. 10.1093/nar/gkw377.

63. Fornes, O., Castro-Mondragon, J., and Khan, A., et al (2020). JASPAR 2020: update of the open-access database of transcription factor binding profiles. Nucleic Acids Res 48(D1):D87-D92.

64. Dreos, R., Ambrosini, G., Groux, R., Cavin Périer, R., and Bucher, P. (2017). The eukaryotic promoter database in its 30th year: focus on non-vertebrate organisms. Nucleic Acids Res 45(D1):D51-D55.

65. Wickham, H. (2016). ggplot2: Elegant Graphics for Data Analysis. Springer International Publishing

66. Wagih, O. (2017). ggseqlogo: a versatile R package for drawing sequence logos. Bioinformatics 33(22):3645-3647

67. Chaudhry, A., Shi, R., and Luciani, D. (2020). A pipeline for multidimensional confocal analysis of mitochondrial morphology, function, and dynamics in pancreatic β-cells. Am J Physiol Endocrinol Metab 318(2), E87–E101

68. Nasir, N., Paus, R., and Ansell, D. (2019). Fluorescent cell tracer dye permits real-time assessment of re-epithelialization in a serum-free ex vivo human skin wound assay. Wound Repair Regen 27(1), 126–133

