## Supplemental Figures for "Mitophagy upregulates WNT5A/Ca^2+^ signalling to accelerate fibroblast migration and wound healing"

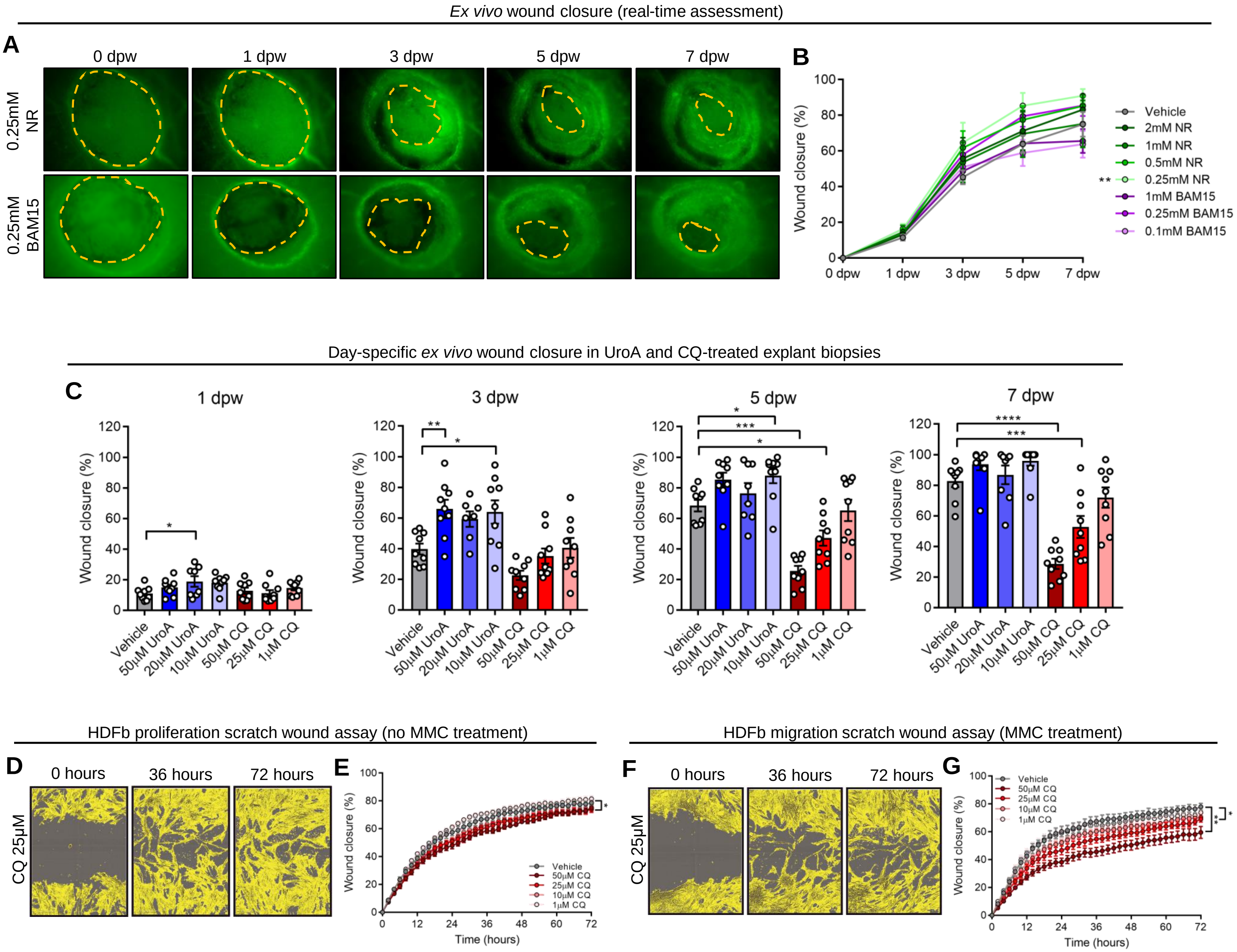


**Figure S1 – Mitophagy accelerates and CQ inhibits re-epethelialisation and HDFb migration, related to Figure 3.**

(**A**) Representative fluorescence images of wounded explant *ex vivo* biopsies treated with vehicle, 0.25mM NR and 0.25mM BAM15 at 0, 1, 3, 5, and 7 dpw. Scale bar = 200μm. (**B**) Quantification (mean ± SEM) of wound healing time of wounds treated with various concentrations of UroA and CQ. Two-way ANOVA,; *** = p < 0.001. N = 5 biological replicates. (**C**) Graphs (mean ± SEM) of wound closure % of vehicle, UroA, and CQ treated *ex vivo* full-thickness explants at 1, 3, 5, and 7 dpw. Two-way ANOVA,; **** = p < 0.0001, *** = p < 0.001, ** = p < 0.01, * = p < 0.05. (**D-E**) Representative images and (**E**) quantification (mean ± SEM) of scratch wound assay in HDFb treated with vehicle and CQ. (**F-G**) Representative images and (**G**) quantification (mean ± SEM) of scratch wound assay in HDFb treated with vehicle + MMC and CQ + MMC over 72 hours. Two-way ANOVA, ** = p < 0.01, * = p < 0.05. N = 3 biological replicates containing at least 3 technical replicates.

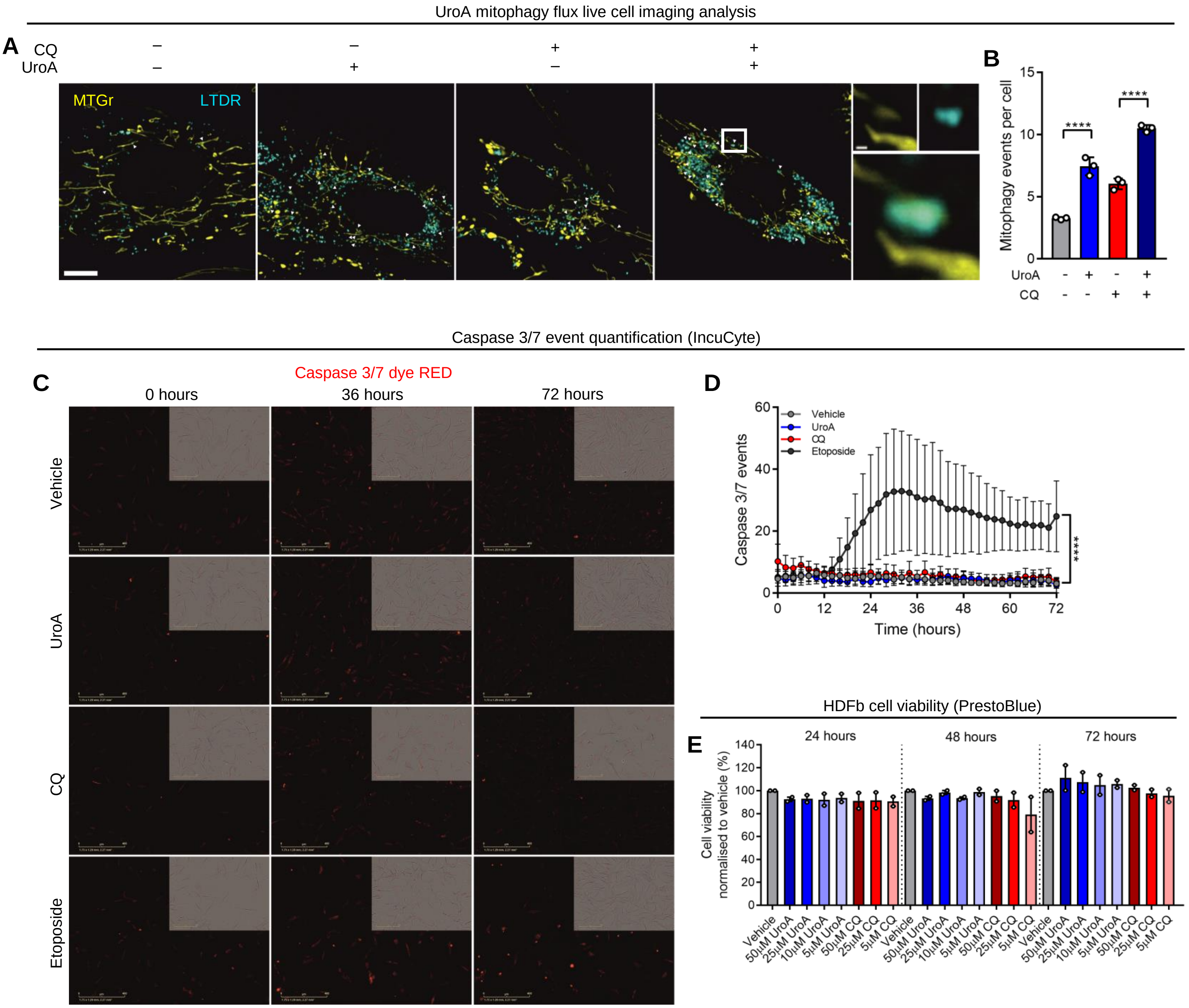


**Figure S2 – UroA increases mitophagy and is not toxic in HDFb, related to Figure 3.**

(**A-B**) Representative fluorescence images and (**B**) quantification (mean ± SEM) of mitophagy events per cell in HDFb following staining with MitoTracker Green and LysoTracker Deep Red. Two-way ANOVA,; **** = p < 0.0001. N = 3 biological repeats containing at least 15 cells. (**C-D**) Representative fluorescence images and (**D**) quantification (mean ± SEM) of Caspase 3/7 events over 72 hours in HDFb stained with Caspase 3/7 dye RED. Two-way ANOVA,; **** = p < 0.0001. N = 3 biological repeats. (**E**) Quantification (mean ± SEM) of percentage cell viability normalised to vehicle in HDFb treated with various concentrations of UroA and CQ over 72 hours.

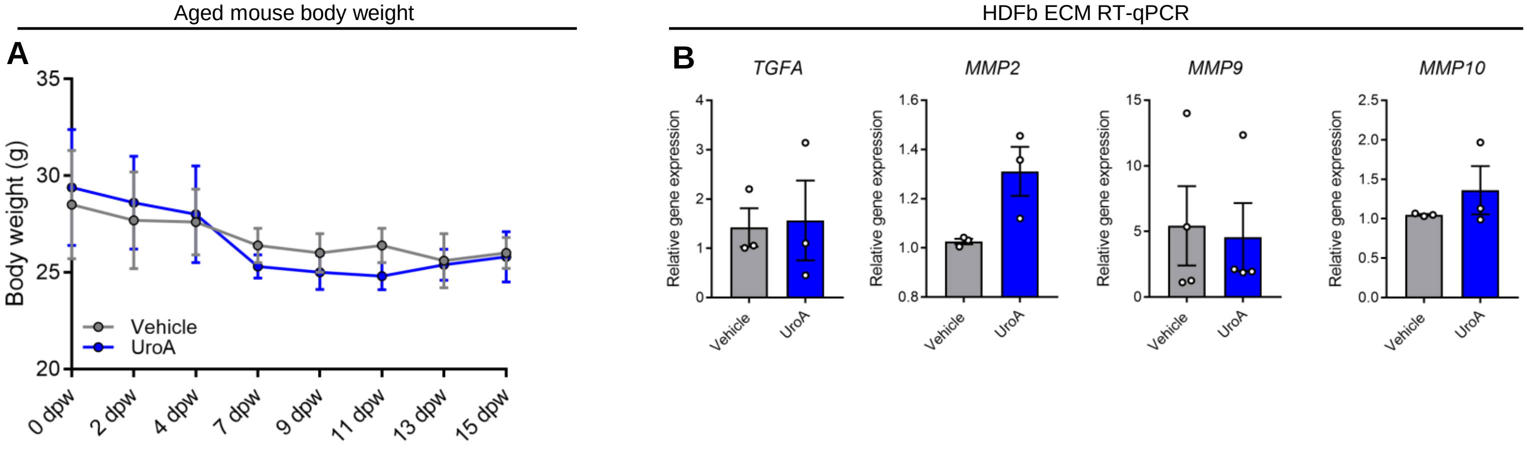


**Figure S3 – No change in mouse body weight and ECM gene expression in HDFb, related to Figure 4.** (**A**) Quantification (mean ± SEM) of body weight (g) of mice treated with either vehicle or UroA for 15 days. Two-way ANOVA. (N = 25 mice). (**B**) Quantification (mean ± SEM) of gene expression of ECM-related related genes in HDFb treated with either vehicle for UroA for 24 hours. Student's t-test. Each dot represents an individual biological repeat containing at least 2 technical repeats.

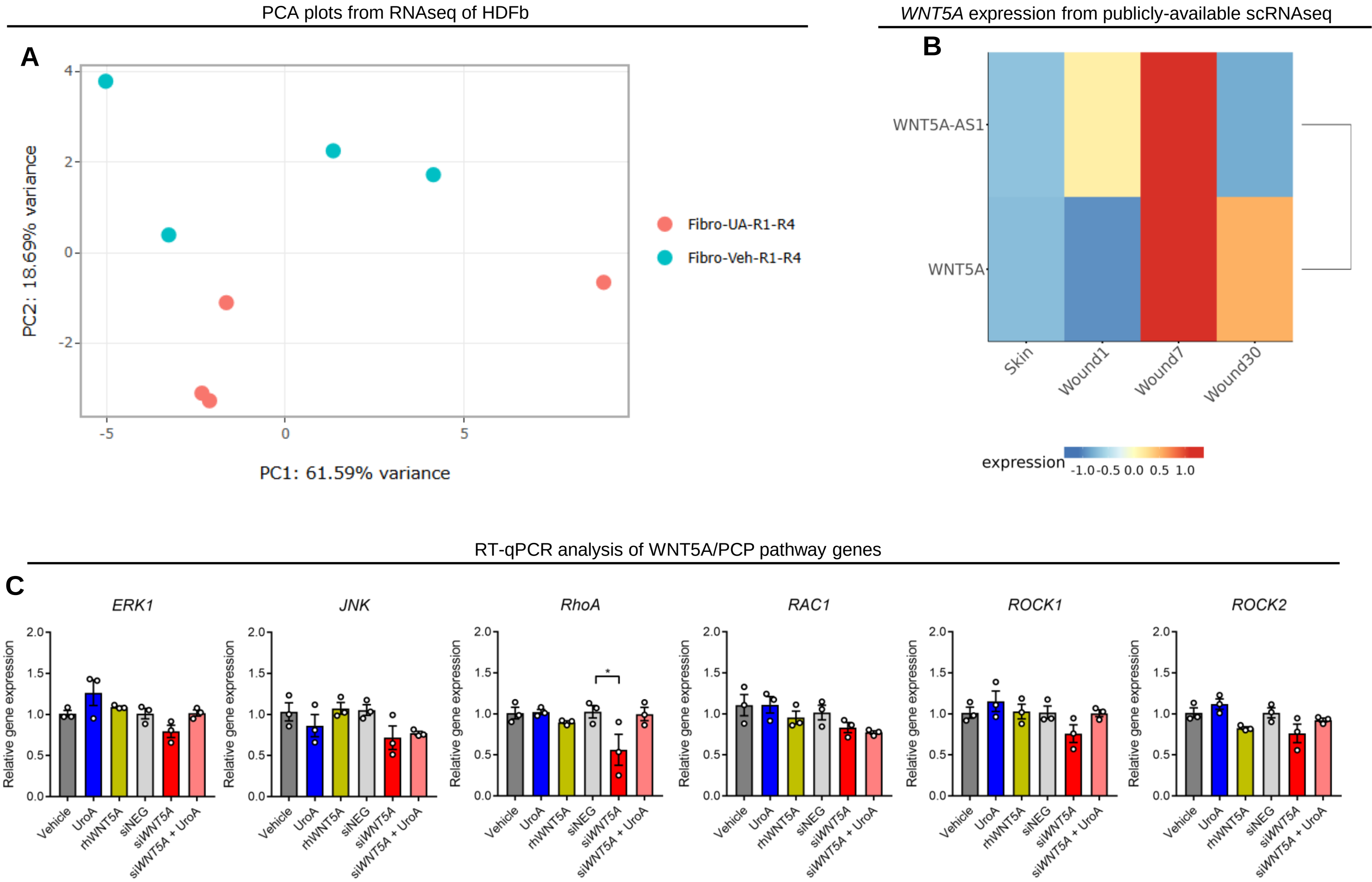


**Figure S4 – *WNT5A* is increased in fibroblasts in human wound healing samples and UroA does not increase WNT5A/PCP pathway genes in HDFb, related to Figure 5.**

(**A**) PCA showing similarity analysis of Vehicle and UroA-treated HDFb used for RNAseq. (**B**) Heatmap showing *WNT5A* and *WNT5A-AS1* expression in FB-1, FB-II, FB-III, and FB-prolif clusters from scRNAseq data, derived from https://shiny.xulandenlab.com/shiny/scwoundatlas/. (**C**) Quantification (mean ± SEM) of the relative gene expression of genes associated with the WNT5A/PCP pathway in HDFb. Two-way ANOVA, * = p < 0.05. N = 3 biological replicates containing at least 2 technical repeats.

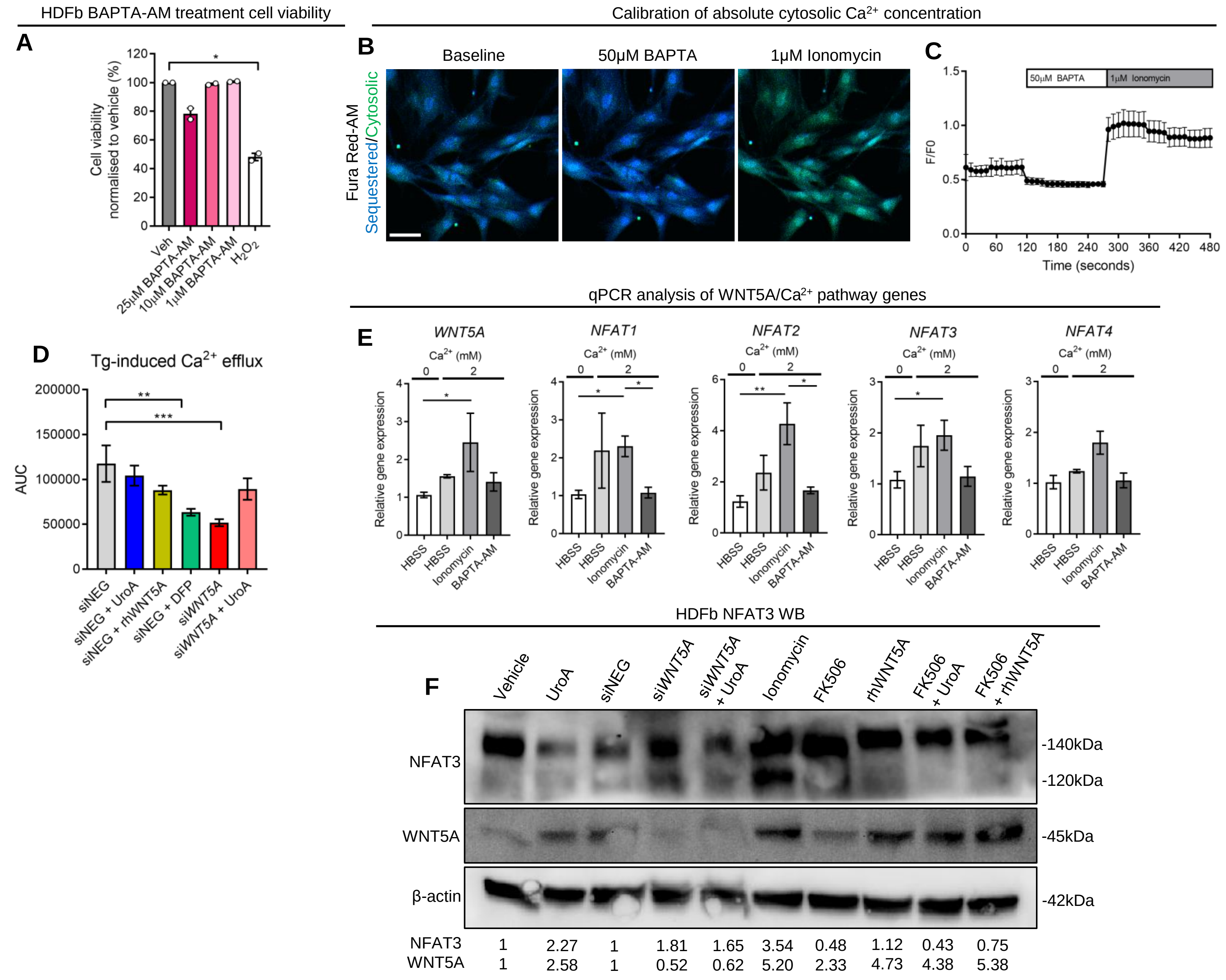


*In sillico* analysis of CREB1 binding sites in *WNT5A*

**G**

| Matrix ID | Name | Score | Relative score | Sequence ID | Start | End | Strand | Predicted sequence |
|---|---|---|---|---|---|---|---|---|
| **MA0018.1** | MA0018.1.CREB1 | 8.742853 | 0.8607433 | FP005546 | 930 | 941 | - | CTGGATGAGGTA |
| **MA0018.1** | MA0018.1.CREB1 | 8.567282 | 0.85602385 | FP005546 | 350 | 361 | - | TGGGTTTACGTA |
| **MA0018.1** | MA0018.1.CREB1 | 7.8948565 | 0.8379487 | FP005546 | 221 | 232 | + | TGGGGGGACGTG |
| **MA0018.1** | MA0018.1.CREB1 | 6.6135707 | 0.80350703 | FP005546 | 344 | 355 | + | CCTGGCTACGTA |

**Figure S5 – $Ca^{2+}$ signalling increases *WNT5A* transcription in HDFb, related to Figure 6.**

(**A**) Quantification (mean ± SEM) of cell viability in HDFb treated with various concentrations of $Ca^{2+}$ quencher BAPTA-AM. Two-way ANOVA, * = $p < 0.05$. Each dot represents an individual biological repeat (N = 2) containing at least 3 technical repeats. (**B-C**) Representative fluorescence images and (**C**) graph of F/F0 ratio of HDFb stained with Fura Red-AM at baseline, after 50µM BAPTA-AM (*Rmin*), and 1µM ionomycin (*Rmax*). (**D**) Quantification (mean ± SEM) of area under the curve after Tg-induced $Ca^{2+}$ efflux in HDFb stained with Fura-Red AM. Two-way ANOVA, *** = $p < 0.001$, ** = $p < 0.01$. N = 3 biological repeats containing at least 20 cells. (**E**) Quantification (mean ± SEM) of *WNT5A* and *NFAT* gene expression in response to $Ca^{2+}$ modulation in HDFb. N = 3 biological repeats containing at least 2 technical repeats. Two-way ANOVA, * = $p < 0.05$. (**F**) Representative immunoblot and relative band expression of NFAT and WNT5A protein levels in HDFb. (**G**) Motif analysis of putative CREB binding sites in the *WNT5A* promoter region.

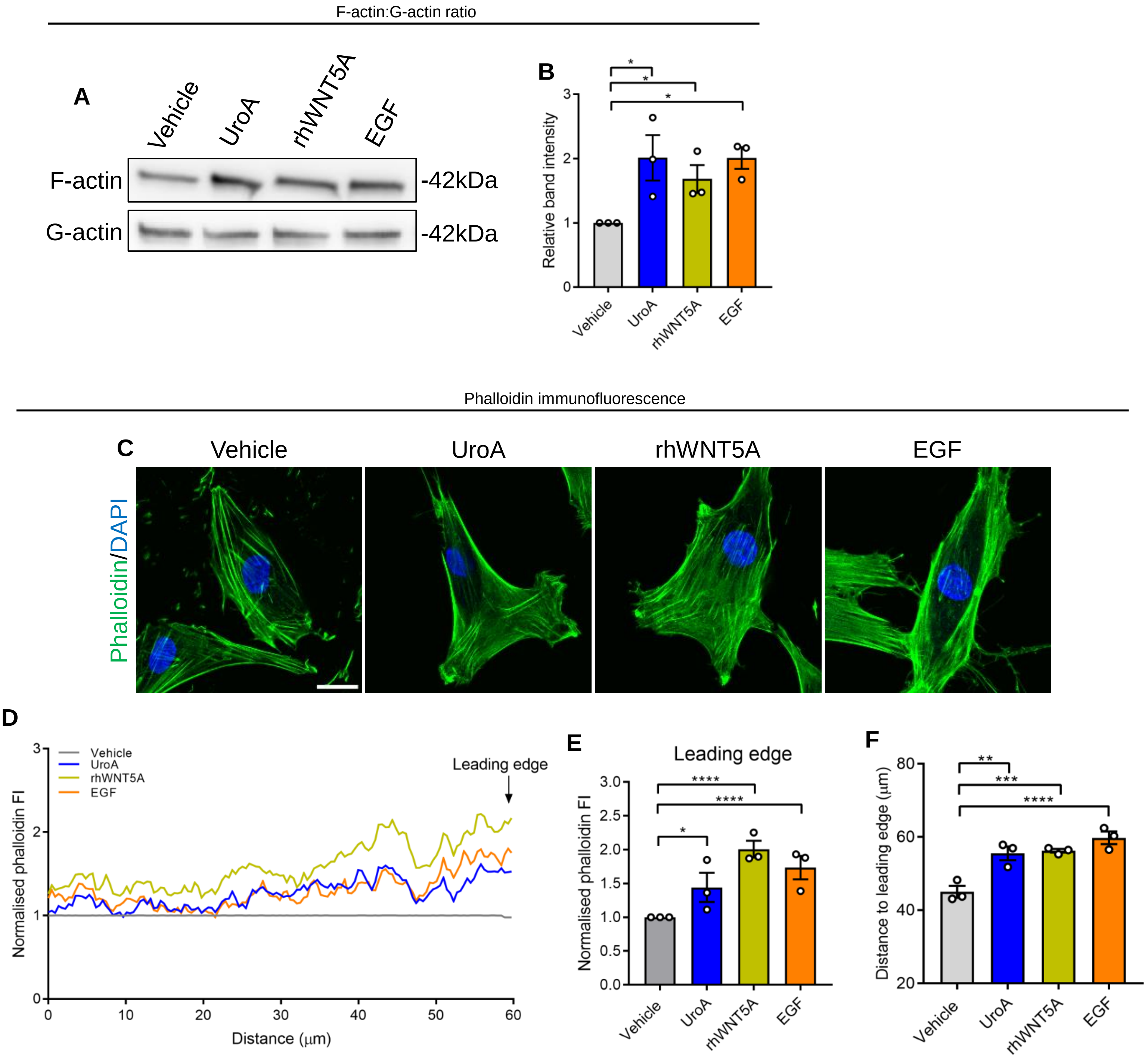


**Figure S6 – UroA, rhWNT5A, and EGF increase actin polymerisation in HDFb, related to Figure 7.** (**A-B**) Representative immunoblot and (**B**) quantification of F-actin and G-actin in HDFb. Each dot represents an biological repeat (n = 3). Two-way ANOVA, * = $p < 0.05$. (**C**) Representative immunofluorescence images of phalloidin staining in HDFb. Scale bar = 10μm. (**D**) Point plot of phalloidin fluorescence intensity normalised to Vehicle in HDFb. Each line represents the mean of 3 biological replicated each containing 15 cells. 60μm indicates the cell periphery. (**E**) Quantification (mean ± SEM) of the average phalloidin fluorescence intensity at the final 10μm of the cell periphery. Two-way ANOVA, **** = $p < 0.0001$, * = $p < 0.05$. Each dot represents an individual biological replicate containing 15 individual cells. (F) Quantification (mean ± SEM) of the distance (μm) from the nuclei to the periphery of the leading edge within HDFb. Two-way ANOVA, **** = $p < 0.0001$, *** = $p < 0.001$, ** = $p < 0.01$. Each dot represents an individual biological replicate containing 15 individual cells.
